# On the Mechanism of Ezrin Activation

**DOI:** 10.1101/2025.11.07.687285

**Authors:** Dovydas Vasiliauskas, Jeriann Beiter, Sahithya Sridharan Iyer, Andrew T. Lombardo, Michelle C. Mendoza, Gregory A. Voth

## Abstract

Ezrin is a peripheral membrane protein that contributes to the organization and stability of cellular membrane structures by reversibly linking the plasma membrane to actin filaments. The formation of this membrane-actin linkage has been experimentally shown to require ezrin N-terminal (FERM) domain binding to PI(4,5)P_2_ phospholipid-enriched membrane sites and the phosphorylation of the ezrin C-terminal domain (CTD) at residue T567. Collectively, membrane association and T567 phosphorylation are believed to promote separation of the FERM and CTD domains; however, the underlying molecular mechanism remains less clear. In this study, we investigate the mechanistic steps of ezrin activation and the thermodynamic free energy landscape of FERM-CTD dissociation using enhanced sampling molecular dynamics (MD). We find that upon ezrin attachment to a lipid membrane, PI(4,5)P_2_ molecules outcompete other phospholipids at the surface of the FERM F1 and F3 subdomains. This interaction triggers a major conformational rearrangement within the FERM domain that destabilizes the FERM F2-CTD interface and initiates dissociation between the FERM and CTD. By employing well-tempered metadynamics (WTMetaD) with a contact-map collective variable, we determine that the principal barrier to FERM-CTD dissociation comes from F3-CTD interactions and that this dissociation can happen spontaneously with a moderate free energy barrier. We also show that the FERM-CTD reassociation after ezrin T567 phosphorylation is impeded due to reduced dissociation energy barrier. The free energy profile of dissociation between FERM and the CTD-replacing EBP50 protein is similar to that of the FERM-CTD system with nonphosphorylated T567, which agrees well with an *in vivo* experimental observation that EBP50 competes with the CTD for F2-F3 binding after CTD is dissociated. Together, our results help establish a revised view on the ezrin activation mechanism where FERM binding to PI(4,5)P_2_ enables spontaneous dissociation of the nonphosphorylated CTD.

**SIGNIFICANCE:** Ezrin and related ERM proteins control how cells link their plasma membrane to the actin cytoskeleton, a process fundamental to cell shape, signaling and motility. Despite decades of study, the molecular basis of ezrin activation – how it transitions from a self-inhibited to an active membrane-bound state – has remained unresolved. Using atomistic and enhanced sampling molecular dynamics together with biochemical validation, we show that binding of the FERM domain to PI(4,5)P_2_-enriched membranes alone is sufficient to trigger spontaneous dissociation of the nonphosphorylated C-terminal domain. Phosphorylation of T567 subsequently stabilizes the open conformation and prevents domain reassociation, enabling actin engagement and binding of FERM partners such as EBP50. Collectively, these findings advance a more integrated view of ezrin activation, highlighting how membrane interactions, conformational flexibility and phosphorylation act in concert to regulate membrane-cytoskeleton coupling.

## INTRODUCTION

Ezrin is a protein of the ezrin-radixin-moesin (ERM) family that is important for the maintenance of actin filament-containing cell surface structures, such as the microvilli, and for participating in cellular signaling responses (1). It is abundant in a wide variety of cultured cells including epithelial tissue cells of intestine, placenta, kidney, lung, stomach and spleen and, if misregulated, can contribute to cancer metastasis (2–6).

Ezrin contains a clover leaf-shaped N-terminal FERM domain (residues 1-296) that consists of three subdomains F1-F3, a long coiled-coil linker domain (residues 297-474) and a C-terminal (CTD) C-ERMAD domain (residues 474-586) (Figure 1a-b). The protein coexists between a closed state where the F2-F3 surface FERM domain is masked by the CTD domain, and an open state where FERM and CTD domains are separated and link the FERM-attached plasma membrane with the CTD-attached actin filaments (7–11). Upon dissociation of CTD from FERM, the F2-F3 surface becomes available for binding of the C-terminal domain of EBP50 protein that participates in cell growth and trafficking pathways (12–14).

**Figure 1.**
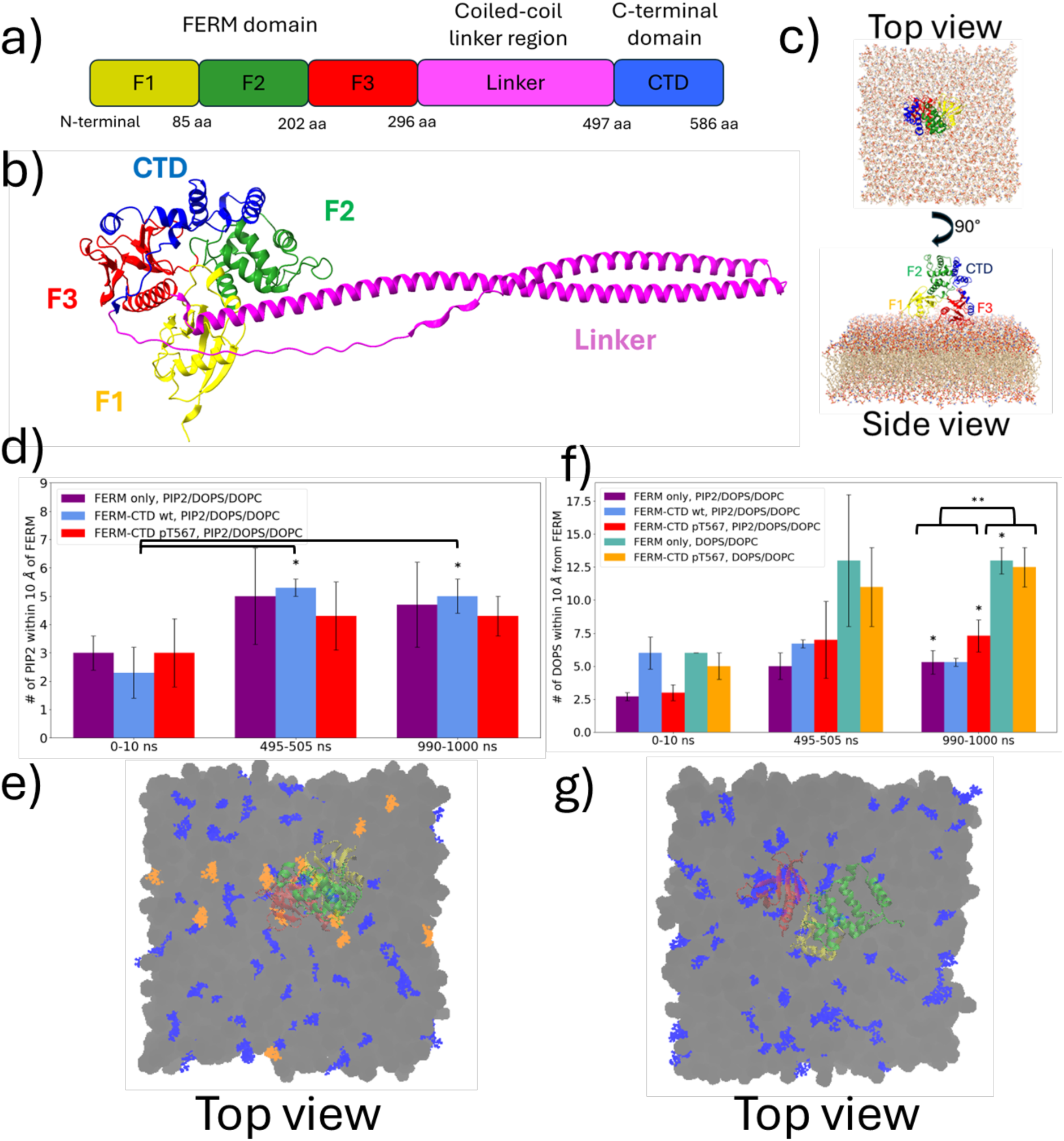
Ezrin FERM domain preferentially binds PIP_2_ over DOPS. a) Ezrin protein sequence: membrane binding FERM domain consists of three lobes – F1 (residues P2-P86), F2 (residues E87-G202), and F3 (residues I203-K296); linker region consists of residues P297-P496; and the actin-binding C-terminal domain (CTD) consists of residues T497-L586. b) AlphaFold 3.0 predicted closed-form structure of the full-length ezrin protein (15). In the predicted closed-form ezrin structure, C-terminal CTD domain interacts with FERM over the F2-F3 subdomain surface. c) System setup for the FERM-CTD structure (PDB ID: 4RM9) at a DOPC:DOPS:PIP_2_ (80:16:4ratio) membrane that was used in the molecular dynamics simulations in this study. d) Number of PIP_2_ head groups phosphorus atoms within 10 Å of any atoms in the residues in FERM. Asterisks (*) indicate significant difference (α = 0.05) between the means of contact counts between 0-10 ns and 495-505 ns or 990-1000 ns, e) Aggregation of PIP_2_ (orange), DOPS (blue) and DOPC (gray) phospholipids in a PIP_2_/DOPS/DOPC membrane at the F1-F3 surface of ezrin FERM domain at *t* = 1000 ns mark. f) Number of DOPS within 10 Å of the residues in FERM. Asterisks (*) indicate significant difference (independent sample t-test, α=0.05) between the means of contact counts between 0-10 ns and 495-505 ns or 990-1000 ns. Double asterisk (**) indicates significant difference (ANOVA, α = 0.05) between the means in systems with DOPC/DOPS/PIP_2_ membranes and DOPC/DOPS membranes in the 990-1000 ns interval. g) Aggregation of DOPS phospholipids (blue) in a DOPS/DOPC membrane at the F1-F3 surface of ezrin FERM domain at *t* = 1000 ns mark.

The dynamics of the ezrin transition from its closed state to open state has been of interest since the early structural studies of ezrin and its ERM family homologs(16). When in closed state, ezrin and other ERM family members have an unusually large (2700 Å^2^) FERM and CTD self-inhibition interface area where the two domains interact through a hydrophobic pocket and hydrogen bonding interactions (16, 17). To overcome the strong FERM/CTD interactions and enter the open membrane-actin tethering state, ezrin undergoes a two-step activation transition. The first activation step requires FERM domain attachment to a PI(4,5)P_2_ (PIP_2_)-enriched membrane plasma membrane through its F1-F3 surface (18–21) and the second step requires phosphorylation of the C-ERMAD T567 residue (21–24). Without phosphorylation of T567 (or a mutation of T567 to a phosphomimetic D567), the CTD domain is inaccessible for actin binding even if PI(4,5)P_2_ binding occurs (25–28).

Recently, computational studies have given more insight into how ezrin FERM domain localizes at PIP_2_-enriched membranes and increases local density of PIP_2_ (29). However, the mechanistic details of how PIP_2_ aggregation and T567 phosphorylation mediate the transition from the closed state to the open state of ezrin remain unknown. In our study, we have investigated the structural changes and energetic profiles during the ezrin activation process using a combination of atomistic molecular dynamics (MD) simulations and well-tempered metadynamics (WT-MetaD) (30, 31) simulations, as well as *in vitro* and *in vivo* cellular experiments. The findings from the MD simulations suggest that ezrin which is bound to a PI(4,5)P_2_-containing membrane and has a phosphorylated T567 residue features weaker interactions between the H1-H2 helices of the CTD domain and the FERM F2 lobe. The results of WT-MetaD simulations with a contact map collective variable give evidence that 1) the unphosphorylated ezrin CTD can spontaneously open and close with a reasonable kinetic barrier and does not require the binding of the kinase LOK to cause this opening (23), and 2) upon phosphorylation of the ezrin CTD residue T567, ezrin is more likely to remain in the FERM-CTD dissociated state due in part to a reduced dissociation barrier. The experimental findings support the *in silico* predictions of free energy profiles by showing that EBP50 attaches to FERM following pT567 phosphorylation and competes with the ERM-CTD for binding at the FERM interface.

## MATERIALS AND METHODS

### Unbiased molecular dynamics simulations

MD simulations were used to investigate different ezrin and membrane systems in aqueous solutions. Structure of the ezrin FERM domain was obtained from PDB 4RMA (containing residues 1-296) and the structure of the closed state ezrin FERM and C-ERMAD system was obtained from PDB 4RM8 (containing residues 1-296 and 519-586) (44). The ezrin linker region was excluded because it was not present in any available crystal structures and the thermodynamics of ezrin activation were expected to be adequately revealed from the crystal structure of the FERM-CTD system. In systems with a membrane and PIP_2_, the membrane was composed of DOPC:DOPS:PIP_2_ phospholipids at a 80:16:4 ratio where the 4% concentration of PIP_2_ is significantly lower than that utilized in previous computational studies of ezrin-membrane interactions (42) and closer to physiological concentrations of PIP_2_ in biological membranes. In systems with membrane that did not contain PIP_2_, a DOPC:DOPS 80:20 composition was used. The lower PIP_2_ concentration was used to minimize the chance of non-specific PIP_2_-FERM interactions and mimic the physiological 1-2% membrane concentration of PIP_2_ as closely as possible. In systems with a membrane, a 15×15 nm^2^ membrane patch was used to fully accommodate the ezrin FERM domain (FERM F1-F3 surface dimensions approx. 3.3×2.5 nm^2^) and leave enough space for lipid regions that do not interact with FERM. Physiological salt concentrations of 0.15 M KCl were used in all simulations. The systems were assembled with CHARMM-GUI molecular simulation preparation tool (45, 46). The simulation were conducted using GROMACS 2022.4 MD software (47–49). Energy minimization was completed using the steepest descent algorithm until the maximum force reached below 1000 kJ/mol/nm. Equilibration and production steps were then carried out in the isothermal-isobaric NPT ensemble using the Nose-Hoover thermostat with time constant τ_t_=1.0 ps, and the exponential relaxation pressure coupling with the Parrinello-Rahman barostat and a reference pressure of 1.0 bar, time constant τ_t_=5.0 ps, compressibility of 4.5×10^-5^ bar^-1^ and semi-isotropic pressure coupling suited for membrane simulations. Production runs used a timestep of 2 fs, the Verlet cutoff scheme, neighbor list updates every 20 steps, Van der Waals cutoff distance of 1.2 nm, particle mesh Ewald method for long-range electrostatic interactions, and hydrogen bond constraints using the LINCS algorithm.

One-microsecond long MD simulations with different combinations of the above-mentioned components were carried out for the following 6 different types of systems: ezrin FERM domain on a membrane with PIP_2_ (3 replicates), ezrin FERM domain on a membrane without PIP_2_ (2 replicates), system of FERM and C-ERMAD with non-phosphorylated T567 and membrane with PIP_2_ (3 replicates), system of FERM and C-ERMAD with phosphorylated T567 and membrane with PIP_2_ (3 replicates), system of FERM and C-ERMAD with non-phosphorylated T567 in solution (2 replicates), system of FERM and C-ERMAD with phosphorylated T567 in solution (2 replicates). A single 500 ns run was conducted for AlphaFold 3.0-constructed complex of ezrin FERM and EBP50 C-terminal domain (residues 225-265) in solution and on a PIP_2_-containing membrane.

The strength of contact between CTD and FERM domain CG beads was evaluated using the heterogeneous elastic network model framework(33). The algorithm allows one to map the CG protein into a network of effective harmonic potentials by iteratively estimating the harmonic spring constants, *k*, based on mean-square distance fluctuations in the pairs of the coarse-grained protein sites.

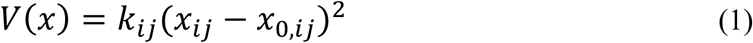

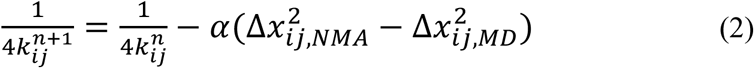

Here *i*, *j* are the respective CG protein sites, 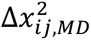 are the mean square distance 4luctuations of the mapped sites of the atomistic molecular dynamics trajectory, and 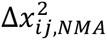 are the projected harmonic fluctuations obtained from a normal mode analysis. Each amino acid residue was mapped to a separate CG bead with the KMC-CG algorithm(50) and the hENM algorithm was implemented using OpenMSCG software(34).

The results were processed, analyzed and depicted using a combination of bash and Python 3.8 scripts.

### Well-tempered metadynamics simulations

The free energy profiles of FERM-CTD dissociation in FERM-CTD systems with phosphorylated and non-phosphorylated T567 at DOPC:DOPS:PIP_2_ membranes were determined through well-tempered contact map metadynamics simulations (30, 31, 51). The contact map collective variable *CMS*(*R*) can be expressed as (52)

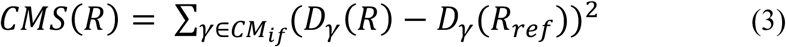

with

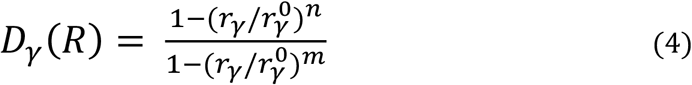

In the context of this study, *CM_if_* was the contact map of Cα residues in the FERM domain that make a shorter than 10 Å contact with the Cα residues of the CTD domain (a total of 230 contacts), *D_γ_*(*R*) is the sigmoid function that defines the contact map contribution of each contact, and *D_γ_*(*R_ref_*) is the sigmoid function values for reference contact pair distances (the contact distance in the starting production frame of the FERM-CTD-npT567 system). For each *D_γ_*(*R*), *r_γ_* is the contact pair distance for any certain WTMetaD simulation frame and 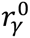 is the contact pair distance in the reference structure. The coefficients *m* = 10 and *n* = 6 make the *D_γ_*(*R*) ≈ 0 if 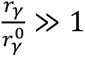 and *D_γ_*(*R*) ≈ 1 if 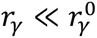. The *CMS*(*R*) values are standardized to range in the [0,1] interval where 1 corresponds to small overall contact distances, or closed system state, and 0 corresponds to large contact distances, or open system state. The WTMetaD simulations were conducted with PLUMED 2.9 software (53, 54) plugged into the GROMACS 2021.5 engine. The contact map WTMetaD was executed using the native contact *Q* switch function implementation (55). Each of the 230 contacts was equally weighted and had beta values of 10.0 and lambda values of 1.2. The Gaussian metadynamics potentials were added with an initial hill height of 1.0 kJ/mol, bias factor of 10.0, and hill width standard deviation of 0.1. Hill additions were executed every 2 ps. Starting frames of metadynamics runs were selected from regularly spaced unbiased MD simulation frames – a total of 32 frames ranging between 250-1000 ns (one every 50 ns) were used for FERM-CTD systems with nonphosphorylated and phosphorylated T567 (16 each), and 8 frames ranging between 150-500 ns (one every 50 ns) were used for the FERM-EBP50 system.

The timescale of transition between the closed and open states of the FERM-CTD system was estimated from the transition state theory derived equation

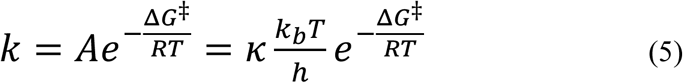

Where *κ* is the transmission coefficient set equal to 1 in this case, *k_B_* and *h* are Boltzmann and Planck constants, respectively. The timescale is calculate as *τ* = 1/*k*.

Docking analysis of ezrin CTD and LOK kinase domain was completed using AlphaFold3 (15) and ClusPro(56) software.

### In Vitro Kinase Assays and Reagents

PI(4,5)P_2_ (cat # 902 Cell Signals, Inc Colombus, OH) were vacuum dried in glass vials and hydrated in kinase buffer (20 mM Tris pH 7.4, 140 mM NaCl, 1 mM EGTA, 1 mM DTT) and sonicated in water bath prior to kinase assay. Kinase assays were performed in kinase buffer with and 200 µM Adenosine 5′-triphosphate (ATP) disodium salt hydrate (Sigma-Aldrich Saint Louis, MO) at 37°C for 10 min. 50 nM of purified kinase was used to phosphorylate 1.5mg/ml of ezrin and 0.5mg/ml of EBP50 or EBP50 variants. PI(4,5)P_2_ was added at 90 µM final concentration. 20% of each kinase assay was boiled in 5X reducing sample buffer in kinase buffer and run on an SDS-page Gel. Western blotting was performed by transferring using a BioRad semi-dry Trans-blot (cat # 1703940) blotting device onto Immobilon-P PVDF Membrane (Millipore cat #IPVH00010). Ezrin mouse monoclonal antibody (DSHB cat# CPTC-Ezrin-1) was used at a dilution of 1:5000 and Phospho-ezrin was detected using rabbit anti-pT567 antibody, raised against recombinant phosphopeptide CRDKYK(pT)LRQIR (57) was used at a dilution of 1:1000. Secondary antibodies used were donkey anti-mouse Alexa Fluor 647 (Thermo Fisher Scientific Cat# A-31571) or IRDye 800CW goat anti-rabbit (LI-COR Biosciences cat# 827–08365). Membranes were imaged using Odyssey CLx imager (Odyssey CLx).

### Protein Expression and Purification

EBP50 constructs were expressed in Rosetta 2(DE3)pLysS cells (EMD Millipore, Billerica, MA). Cells were grown in Terrific broth with antibiotics at 37°C up to OD 1.0. Proteins were expressed at 30°C for 4 hr by adding 1 mM IPTG. EBP50-SUMO proteins were lysed from cells in binding buffer (20 mM sodium phosphate, 300 mM NaCl, 20 mM Imidazole, pH 7.4), cleared and filtered prior to binding to Ni-NTA agarose (Qiagen, Hilden, Germany) for 2 hr. at 4C. SUMO fusion protein was eluted (20 mM sodium phosphate, 300 mM NaCl, 500 mM Imidazole, pH 7.4) and dialyzed against binding buffer. EBP50 construct’s 6xHis-SUMO tag was cleaved with 6×His-ULP1 protease in presence of 1 mM DTT and removed with Ni-NTA. Ezrin and ezrin FERM were expressed in M15 cells (Qiagen, Hilden, Germany) at 27°C for 4.5 hr, then lysed in Buffer A (180 mM KH2PO4, 180 mM K2HPO4) and purified over a hydroxyapatite column by gradient elution(12). LOK-GFP-flag was isolated by flag pulldown from HEK 293T (ATCC Cat# CRL-1573) cells stably expressing the plasmid (23). In brief cells were washed with phosphate-buffered saline (PBS) lysis buffer (25 mM Tris, 5% glycerol, 150 mM NaCl, 50 mM NaF, 0.1 mM Na3VO4, 10mM βGP, 0.2% Triton X-100, 250 mM calyculin A, 1 mM DTT, 1× cOmplete Protease Inhibitor Cocktail [Roche; Cat# 11836153001]) was added. Lysates were then sonicated, and centrifuged at 8000 × g for 10 min at 4°C. The supernatant was taken and added to M2-Flag beads (sigma cat# A2220) and nutated for 3 hr at 4°C. Protein was eluted with 3x flag peptide (sigma cat# F4799).

### Cell Culture

Jeg-3 cells (ATCC Cat# HTB-36) and HEK 293T (ATCC Cat# CRL-1573) were maintained in 5% CO2 humidity at 37°C in MEM (Life Technologies, Waltham, MA) with 10% FBS (Life Technologies, Waltham, MA) and 1X GlutaMAX (Life Technologies, Waltham, MA). CRISPR knockout cell lines were produced and validated earlier for LOK and SLK (58) and EBP50 (41). Cells were treated with Calyculin A (Cell Signaling technologies cat #9902) at 1µM for 5min or Staurosporine (Sigma-Aldrich cat# AM2282) at 1µM for 5min. For analysis of endogenous ezrin phosphorylation, cells were rapidly boiled in 5X reducing sample buffer in kinase buffer, diluted 3-fold, vortexed vigorously, and cleared by centrifugation prior to SDS-PAGE before western blotting.

## RESULTS

To understand specific effects of ezrin attachment to a PIP_2_-enriched membrane and phosphorylation of the CTD residue T567, simulations were conducted for three different types of ezrin systems – FERM only domain (ezrin residues 2-296, PDB structure 4RMA), as well as FERM-CTD constructs (ezrin residues 2-296 and 519-586, PDB structure 4RM9, Figure 1c) with phosphorylated T567 and FERM-CTD construct with nonphosphorylated T567. The evolution of each of the system was tested in three different environments – either a DOPC:DOPS:PIP_2_ (80:16:4 ratio) membrane, a DOPC:DOPS (80:20 ratio) membrane, or in the absence of a membrane (solution-only). Microsecond-long simulation replicas were collected for each of the different systems as explained in the Methods section. The results below are averaged over the independent replicate runs and presented with the standard error bars. Full length ezrin (Figure 1b) was not considered in order to reduce the complexity added by the linker region and to facilitate the analysis of the statistical mechanics of FERM-CTD dissociation.

### PI(4,5)P_2_ enables FERM/membrane binding and allosteric modulation of an F2 domain motion

Unbiased MD simulation results indicate that upon binding to a DOPC:DOPS:PIP_2_ or DOPC:DOPS membrane, the FERM domain remains localized and does not dissociate from the membrane throughout the course of the 1 µs simulation (Supplementary Figure S1). The vertical projected z-distance distance between the backbone of the FERM residues that are closest to the membrane head groups (residue groups 2-4, 21-28, 60-65 and 250-260) remains virtually unchanged during the simulations. Upon binding to the DOPC:DOPS:PIP_2_ membrane, the membrane-facing lysine and arginine residues make stable contacts with the PIP_2_ head groups and can recruit additional PIP_2_ head groups during the 1 µs simulation (Figure 1d,1e, Supplementary Figure S2). The number of PIP_2_ head groups within 1 nm of FERM increase from about 3 the start of the simulation to about 5 at the end the 1 µs simulation (Figure 1d,1e). The residues with the most pronounced PIP_2_ head group contacts are R40, K65, K262, R275, K278, R293 and R295 (Supplementary Figure S2). The PIP_2_ head groups attach to the surface of FERM domain lobes F1 and F3 as well as at the cleft between them (Supplementary Figure S3), as expected from previous structural studies(29, 32). In addition to PIP_2_, DOPS (another negatively charged lipid), is sequestered around the FERM domain.

When PIP_2_ is absent from the lipid membranes, the FERM F1 and F3 lobes again recruit DOPS lipids, resulting in an increase from about 5 to about 13 stable contacts with DOPS head groups throughout the course of the 1 µs simulation (Figure 1f,1g). However, when present, PIP_2_ outcompetes DOPS in forming contacts with the positively charged residues in the FERM F1 and F3 lobes as the DOPC/DOPS/PIP_2_ membranes had significantly fewer DOPS contacts with FERM than the DOPC/DOPS membranes (Figure 1f). This is consistent with work *in vivo* that has indicated the importance of PIP_2_ for ezrin recruitment to specific regions of the plasma membrane, and emphasizes the need of PIP_2_ in creating a strong electrostatic attachment(21, 23).

The presence of PIP_2_ translates into structural effects on the FERM domain as well, promoting ezrin activation. The FERM domain on a PIP_2_/DOPS/DOPC membrane was found to have a significant shift in distances between specific secondary structures of the F3 subdomain with respect to F1 and F2 subdomains when compared to other analyzed systems FERM systems on a DOPC/DOPS membrane or in aqueous solution (Figure 2). Most noticeable internal distance changes in this were found for the FERM F3 lobe β-sheet residue group G202-K211 which moved closer to residue groups L76-A82, P119-F134 in F1 and F2 subdomains, as well as the F3 residue group residue group K254-K262 which moves away from the F1 lobe residues P86-E92 and approaches other F3 residue groups G224-D232 (Figure 2a). The FERM systems at the membrane also had higher root mean square fluctuation (RMSF) values than the FERM system in solution in the second half of the 1 µs MD simulations (especially in the F2 residues D136-L152) which further supports the observation that FERM interactions with negatively charged phospholipid head groups induce more motion across the entire FERM domain (Figure 2b).

**Figure 2.**
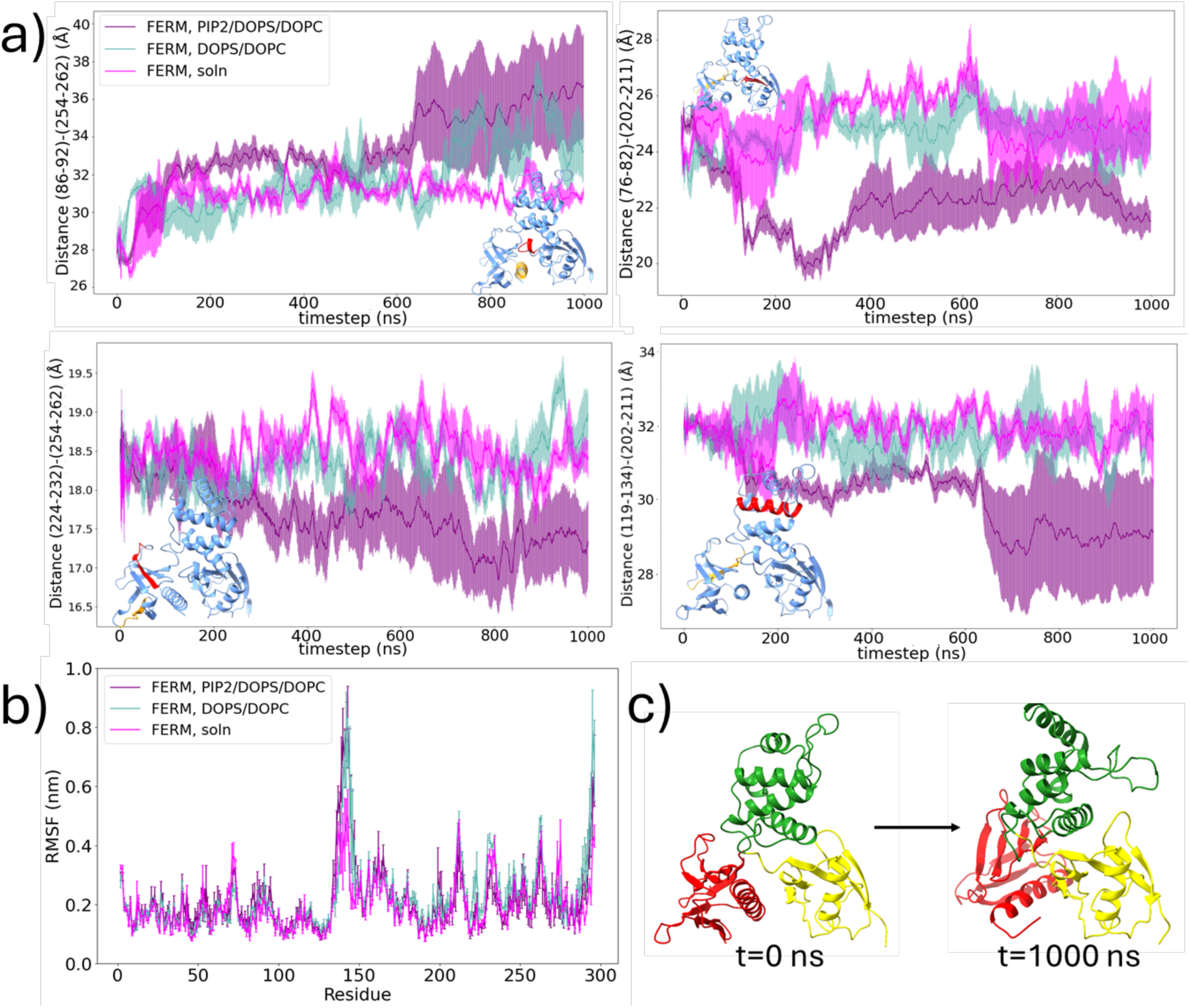
Ezrin FERM domain is activated by PIP_2_ binding. (a) Time series of center-of-mass distance of sample FERM secondary structure pairs. The secondary structures undergo significantly larger conformational change at a PIP_2_/DOPC/DOPS membrane (purple) than at DOPC/DOPS membrane (teal) or solution (magenta). The data is represented as a mean distance of the secondary with standard error across independent replicates of the simulation runs. The figure insets depict the structure of FERM with the specific secondary structures highlighted in red and orange. (b) FERM root mean square fluctuation results. (c) FERM conformational change at a PIP_2_-containing membrane.

Most significantly, the FERM bound to the PIP_2_-containing membrane underwent a rearrangement of the F2 and F3 lobes with respect to the F1 lobe (Figure 2c). This domain-opening motion shows that the F2 domain is quite flexibly linked to the other two lobes, and that anchoring of the F1 and F3 lobes to the membrane via PIP_2_ results in an allosteric propagation of that effect. This motion is significant because it perturbs the ezrin CTD binding interface across F2 and F3 and suggests that F2 domain flexibility is important for initiating CTD dissociation.

### Effects of T567 phosphorylation on FERM-CTD association strength

To examine the effects of threonine 567 (T567) phosphorylation, we incorporated ezrin CTD into our atomistic MD simulations of the FERM domain bound to a PIP_2_-containing membrane. The FERM-CTD system with phosphorylated T567 at the PIP_2_-containing membrane breaks significantly more hydrogen bonds between the FERM F2 subdomain and the CTD than a respective FERM-CTD system with nonphosphorylated T567 (Figure 3a). Residues D136-L152 of F2 fluctuate significantly more in FERM-CTD systems with phosphorylated T567 that are attached to PIP_2_/DOPS/DOPC membrane than in the other analyzed systems (Supplementary Figure S4a-b). The high flexibility of residue group D136-L152 might be expected because these residues are part of disordered loop and are distant both from the motion-restricting membrane as well as the CTD domain (Supplementary Figure S4c). Consistent with the observations of F2 domain motion, these findings suggest that PIP_2_ binding and phosphorylation of T567 cooperatively help to weaken the F2-CTD hydrogen-bonding interaction network and in this way help initiate the dissociation of CTD domain at the F2-CTD surface.

**Figure 3.**
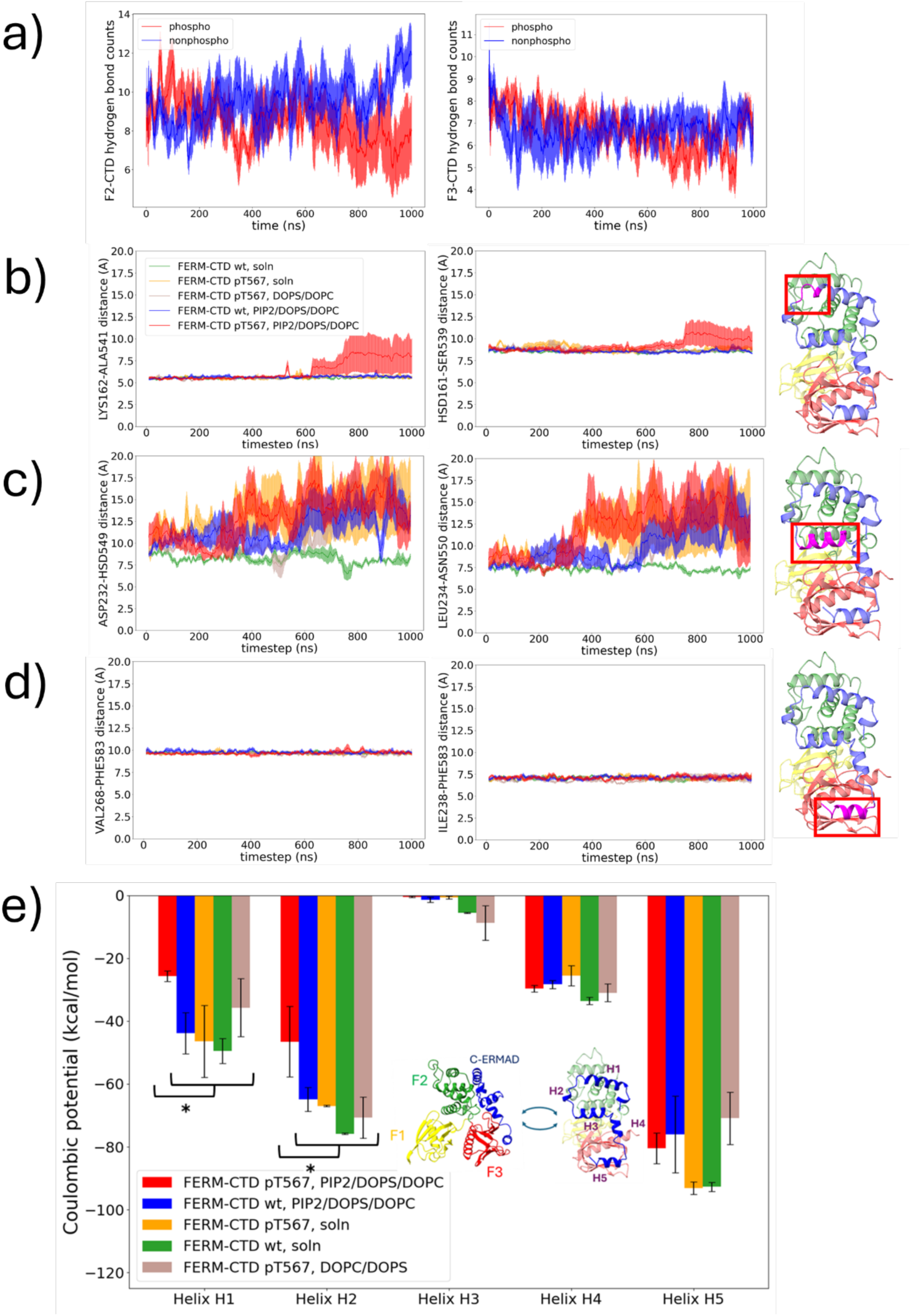
Differential electrostatic interactions throughout the helices of the CTD drive allosteric rearrangements upon phosphorylation. a) Hydrogen bond counts between ezrin F2-CTD lobes (left) and ezrin F3-CTD lobes (right). Significantly more hydrogen bonds are broken in the system where ezrin has a phosphorylated T567 residue. b) Significant opening between CTD residues AA538-AA546 and FERM F2 lobe is only observed in the FERM-CTD system with phosphorylated T567 at a PIP_2_/DOPS/DOPC membrane. Left – crystal structure of ezrin FERM and C-ERMAD (PDB ID:4RM9). Right – distance between alpha carbon atoms of residue pairs K162-R542, Q160-K546, R156-T548 in the beginning of the equilibrated unbiased MD simulations (t=0 ns) and the end of the 1 µs simulation (t=1000 ns) for versions of the system that contain either a nonphosphorylated T567 (npT567, top) or phosphorylated T567 (pT567, bottom). b)-d) Distances of CTD residue Cα atoms from the Cα atoms of their nearest FERM F2-F3 residues. b) distances of CTD residues AA538-AA542 increase only for a system phosphorylated T567 at a PIP_2_/DOPS/DOPC membrane (red), c) large changes in distances are observed between Cα atoms of CTD helix H3 (residues AA549-AA559) and FERM F3 residues K230-T235 an CTD helix H3, d) no significant change in distance is observed between CTD helix H5 (residues T576-L586) and nearest FERM atoms. e) Coulombic potentials between CTD helices H1-H5 and nearest secondary structures in FERM F2-F3 lobes. Helices H1-H5 correspond to amino acid residues as follows: H1 – residues E525-Q540, H2 – residues A541-R547, H3 – residues H549-R559, H4 – residues K564-R572, H5 – residues T576-L586. Notation: pT567 – system with a phosphorylated T567, wt – system with a nonphosphorylated T567, soln – simulation carried out in a KCl solution. Asterisks (*) indicate a statistically significant coulombic attraction strength difference for a certain CTD helix between the FERM-CTD pT567 system at a PIP_2_/DOPC/DOPS membrane against any other systems. The inset provides a visualization of H1-H5 CTD helices in the crystal structure of closed state ezrin (PDB ID: 4RM9).

The compounded effect of T567 phosphorylation and PIP_2_ binding in initiating the F2-CTD dissociation is noticeable from the analysis of distances of FERM-CTD residue pairs that have Cα distances of less than 1 nm (Figure 3b-d). The FERM-CTD pair distances increase for CTD H3 helix of residues H549-R559 (Figure 3c) and do not increase at all for CTD H5 helix of residues T576-L586 in all systems (Figure 3d): this agrees with the CTD RMSF plots that show high flexibility of CTD helix H3 and low flexibility of CTD helix H5 (Supplementary Figure S5). However, a segment of residues E537-R542 between CTD helices H1 and H2 feature an increase of distance from FERM F2 residues only for the system of FERM-CTD with phosphorylated T567 at a DOPC/ PIP_2_/DOPS membrane (Figure 3b). As can be seen in Supplementary Figure S6, the FERM-CTDpT567 system at the DOPC/ PIP_2_/DOPS membrane is the only system that is able to have a substantial separation of the CTD H1 helix from the FERM F2 helix during the course of 1 µs of unbiased MD simulations.

The Coulombic FERM-CTD potentials (Figure 3e) also suggest that interactions at the F2-CTD interface are weaker than at the F3-CTD interface. Helices H1 and H2 have significantly weaker interactions with FERM structures in the system FERM-CTDpT567 at PIP_2_/DOPS/DOPC membrane than in other systems (–25.6 kcal/mol against –36 to –50 kcal/mol in H1 helix and –46 kcal/mol against –64 to –76 kcal/mol in H2 helix). Such significant drop in coulombic interactions is assumed to be a consequence of the separation between H1-H2 helices of CTD from FERM F2 subdomain. The strongest coulombic interactions were observed between CTD helix H5 and the FERM structures (–70 to –95 kcal/mol) which again agrees with the findings that distances of FERM-CTD contacts that involved H5 helices do not change at all (Figure 3e). The CTD residue group H4 that involved the T567 residue had moderate Coulombic interactions with the FERM structures (–24 to –31 kcal/mol) and were not different between the systems that had phosphorylated or non-phosphorylated T567. It is therefore suggested that the insertion of a bulky, negatively charged phosphate group in phosphorylation of T567 reduces the attractive FERM-CTD interactions by inducing a shift in the salt bridge network across the entire surface of the CTD domain.

More insight into the T567-induced salt bridge network shift across CTD comes from heterogeneous elastic network (hENM) model(33). To generate the hENM model, we mapped each amino acid of the FERM/CTD complex to a single CG site and then evaluated the network for bead pairs within 10 Å radius of each other in the 500-600 ns and 900-1000 ns intervals of the atomistic trajectories (Figure 4a-b) (34). It can be observed that the nonphosphorylated FERM-wtCTD had higher estimated harmonic spring constant (*k*) for F2-bordering CTD residue groups Q530-Q540 (CTD helix H1) and F3-bordering groups Y565-Q570 (helix H4) and I580-L586 (helix H5) than the phosphorylated FERM-CTDpT567 system. As presented in the histogram of *k* across all FERM-CTD pairs (Figure 4c), the nonphosphorylated system has statistically significantly stronger FERM-CTD interactions than the system with phosphorylated T567 (Mann-Whitney P=0.031). Taken together, these results suggest that electrostatic repulsions of pT567 weaken the CTD-FERM interactions across the binding interface, rather than locally around just T567.

**Figure 4.**
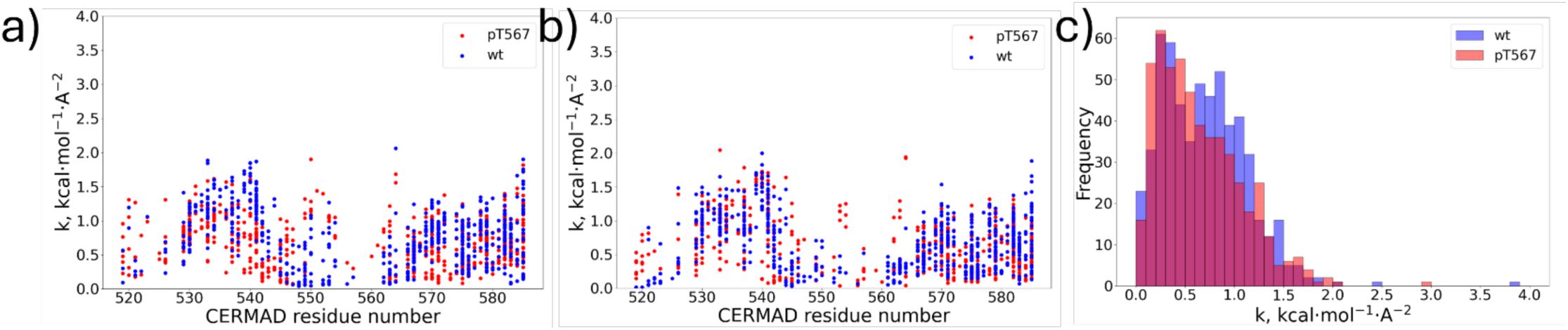
Heterogeneous elastic network modeling (hENM) indicates that CTD helices H2, H4 and H5 have a weaker association with FERM when phosphorylated (red). a)-b) *k* values for each of the CTD residue coarse-grained (CG) sites (AA519-AA586) with all FERM CG sites that are within 15 Å from the CTD site across three independent replicates of simulation sets. a) *k* values for CTD CG residue sites mapped from the atomistic unbiased MD trajectory between 500-600 ns. b) *k* values for the CTD CG residue sites mapped from unbiased MD trajectory between 900-1000 ns. c) frequencies of *k* values of hENM harmonic potentials in the phosphorylated (red) and nonphosphorylated (blue) system across the entire set of FERM-CTD pairs. The distributions are statistically significant (Mann-Whitney U test P=0.031).

### Examination of FERM-CTD dissociation thermodynamics profile using well-tempered metadynamics (WTMetaD) contact map collective variable

To further explore the full dissociation of the CTD from the membrane-bound FERM domain, we explore the thermodynamics of the process via enhanced sampling. WTMetaD (31, 35) ( 2) was well-suited for our system as it does not require a prior knowledge of the intermediate states and employs a history-dependent bias which allows efficient barrier crossing in systems which have a considerable dissociation barrier. To effectively implement WTMetaD the ideal collective variable (CV) must be able to account for the predicted cooperative mechanism of CTD association/dissociation to FERM(36) resulting from the previously described network of salt bridges between F2-F3 subdomains and CTD.

Using a contact map CV that included all pairs of F2-F3 and CTD residues the Cα atoms of which had a contact of less than 10 Å (a total of 230 contacts in each of the FERM-CTD simulations), we were able to effectively sample the full FERM-CTD dissociation. The system was able to sample both the closed system state where FERM-CTD were associated with each other (cmap values of ∼ 0.5 to 1.0) and the open system state where FERM-CTD had dissociated and lost intermolecular contacts (cmap values of 0 to ∼ 0.5). Multiple replicas of WTMetaD runs with starting conformations from equally spaced points of the 1 µs unbiased MD simulations were collected for FERM-CTD systems with phosphorylated and non-phosphorylated (WT) T567 at DOPC/DOPS/PIP_2_ membrane to probe the effect of T567 phosphorylation. With WTMetaD, the system sampled across the entire surface of F2-F3 and CTD leading to full CTD dissociation in 10-20 ns (Figure 5a).

**Figure 5.**
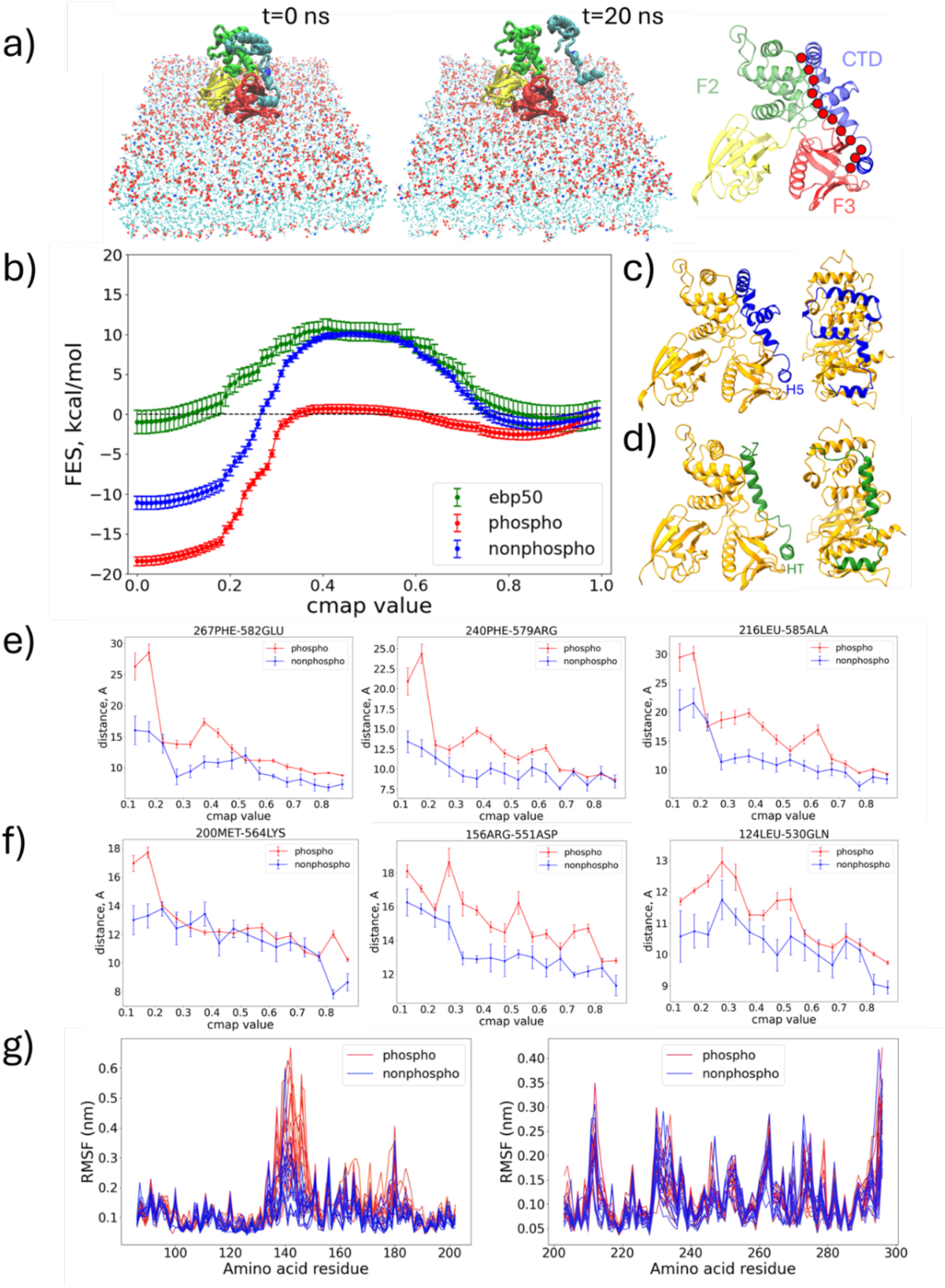
Phosphorylated ezrin FERM-CTDp567 system has a significantly lower FERM-CTD dissociation barrier than the nonphosphorylated system. a) Well-tempered metadynamics contact map collective variable can be successfully used to make the ezrin system undergo a full dissociation between FERM and CTD domains in about 20 ns. Image on the right is the schematic of how the contact map CV includes contacts across the entire surface of CTD and FERM F2-F3 lobes. b) The free energy surfaces of well-tempered contact map WTMetaD show that ezrin system with nonphosphorylated T567 has a mean barrier of 11.2 ± 0.3 kcal/mol for the transition from closed to dissociated FERM-CTD (blue) and a 3.2 ± 0.5 kcal/mol transition barrier between the closed and open states when the system has a phosphorylated T567 residue. The barrier of EBP50-FERM dissociation (11.7 ± 1.1 kcal/mol) is similar to that of FERM-wtCTD. Results are reported as mean (± SE) of sample free energy values corresponding to certain contact map collective variable values. The free energy profiles converged in 50-100 ns of the WTMetaD runs (Supplementary Figure S8). c) Ezrin FERM-CTD crystal structure (PDB ID: 4RM9). d) Crystal structure of ezrin FERM domain and EBP50 C-terminal residues (PDB ID: 1SGH). The C-terminal end of ezrin CTD domain has very similar alignment to F3 lobe of FERM as does the end helix of EBP50. e)-f) Distance between the C_α_ atoms of different contact map pairs at contact map values 0.1-0.9 obtained from WTMetaD runs between F3 and CTD (e), and between F2 and CTD (f). Results in e)-f) are presented as mean distance and standard error for each of the contact map coordinate values collected from all WTMetaD simulation frames that have a contact map value in a certain range. g) RMSF plots for lobe F2 (left) and F3 (right) during the WTMetaD simulations for the system with phosphorylated T567 (red) and the system with nonphosphorylated T567 (blue).

The difference of the free energy landscape in FERM-CTD interactions with phosphorylated and non-phosphorylated T567 was evaluated from the averaged free energy surfaces of each of the conducted WTMetaD runs (Figure 5b). The free energy barriers to dissociation were 11.2 ± 0.3 kcal/mol for the wtCTD system and 3.2 ± 0.5 kcal/mol for the pT567CTD system. Additionally, the barriers to reassociation in wtCTD (21.0 kcal/mol) and pT567 CTD (19.4 kcal/mol) were similar, suggesting that the dissociated FERM-CTD state that favors the pT567 ezrin is primarily facilitated by the high reassociation barrier. The results also suggest wtCTD can kinetically fluctuate from closed to open state without the action of a kinase to first “force it open” before the phosphorylation reaction. These findings also hint that that a main role of pT567 phosphorylation could be to further drive this conformational change to expose the actin-binding interface of the CTD. It is worth noting that the free energy curves (Supplementary Figure 7) for ezrin in solution (not bound to the membrane) also show the ability to fluctuate to an open state so that the phosphorylation of T567 by LOK can still occur(37–39). The kinetic barrier is higher than in the membrane-bound case for the wtCTD by about 3 kcal/mol and, importantly, the free energy for the process is uphill. The solution-phase systems also feature a lower free energy barrier of reassociation than systems at the membrane, implying that membrane further facilitates ezrin transition to open state by disfavoring FERM-CTD reattachment.

To gain additional insight into FERM F2/F3 binding interface, we measured the free energy of the FERM domain interacting with the CTD of ezrin-radixin-moesin-binding phosphoprotein 50 (EBP50, PDB ID: 1SGH). EBP50-CTD (40) shares the same secondary structure and orientation as the FERM-CTD construct when binding to the ezrin FERM domain (Figure 5c-d). The dissociation free energy profile of FERM-EBP50 system at the PIP_2_/DOPS/DOPC membrane was similar to that of the wtCTD system, with a dissociation barrier of 11.7 ± 1.1 kcal/mol and a reassociation barrier of 11.8 kcal/mol (Figure 5b). This suggests that EBP50-CTD is able to bind opportunistically to FERM after FERM-CTD dissociation, thereby prolonging CTD exposure to kinase phosphorylation.

The structural analysis of the FERM-CTD systems from contact map WTMetaD runs supports the results of unbiased MD simulations. The RMSF profiles of FERM domain in the independent WTMetaD runs show that FERM domain had significantly higher mobility and flexibility of the F2 subdomain residues 136-152 in the FERM-CTDpT567 system than the FERM-wtCTD system (Figure 5e). The WTMetaD distance profiles of FERM-CTD contacts across the contact map CV coordinate indicate that the FERM-CTDpT567 systems also had a significantly larger increase in distances of pairs that involved atoms from CTD helix H5 (Figure 5f) and somewhat larger increase in distances for pairs involving atoms of helices H1-H4 (Figure 5g). This once again suggests that the CTD domain dissociates more easily and allows more freedom of motion in the FERM F2 subdomain in the FERM-CTDpT567 system than in the FERM-wtCTD system.

### Experimental validation of *in silico* predictions

The present results predict that phosphorylation at T567 must occur before EBP50 binding to the FERM can occur. This is because the FERM-CTDpT567 system is the only condition where the energy landscape promotes full detachment of the CTD from FERM, subsequently allowing the EBP50-FERM binding domain access to the F2-F3 surface. An alternative hypothesis previously proposed that PIP_2_ binding to FERM partially opens the CTD-FERM interface enabling FERM binding proteins, such as EBP50, to “wedge” opens the FERM-CTD interface followed by stabilization through phosphorylation at pT567. To test between these two hypotheses and if our *in silico* models were predictive, we performed biochemical kinase assays to test if EBP50 enhanced phosphorylation at Ezrin’s T567 residue *in vitro* and in cells. We purified a tagged construct of the endogenous human kinase for ERM’s T567 residue, Lymphocyte Oriented Kinase (LOK), using the HEK-293T expression system (Figure 6a). We incubated LOK with purified full length bacterially expressed ezrin in the presence and absence of PI(4,5)P_2_ and measured the level of pT567 phosphorylation using a phosphospecific antibody compared to total ezrin expression (Figure 6b). As previously reported (23), LOK phosphorylated Ezrin in the presence of PI(4,5)P_2_ but achieved little phosphorylation in the absence of PI(4,5)P_2_. We then tested if the addition of EBP50 to the kinase assay enhanced the levels of pT567 produced in the presence or absence of PI(4,5)P_2_ and found that EBP50 did not affect the phosphorylation of Ezrin’s T567 residue. EBP50’s FERM binding motif is regulated by two PDZ binding motifs within EBP50 and mutation of these motifs or truncation to include only the final 38 amino acids of the EBP50 tail produce a constitutively active FERM binding EBP50 constructs. These EBP50 variants also did not affect phosphorylation at the T567 residue under any conditions. We concluded that EBP50 does not promote the phosphorylation of Ezrin at the T567 residue.

**Figure 6.**
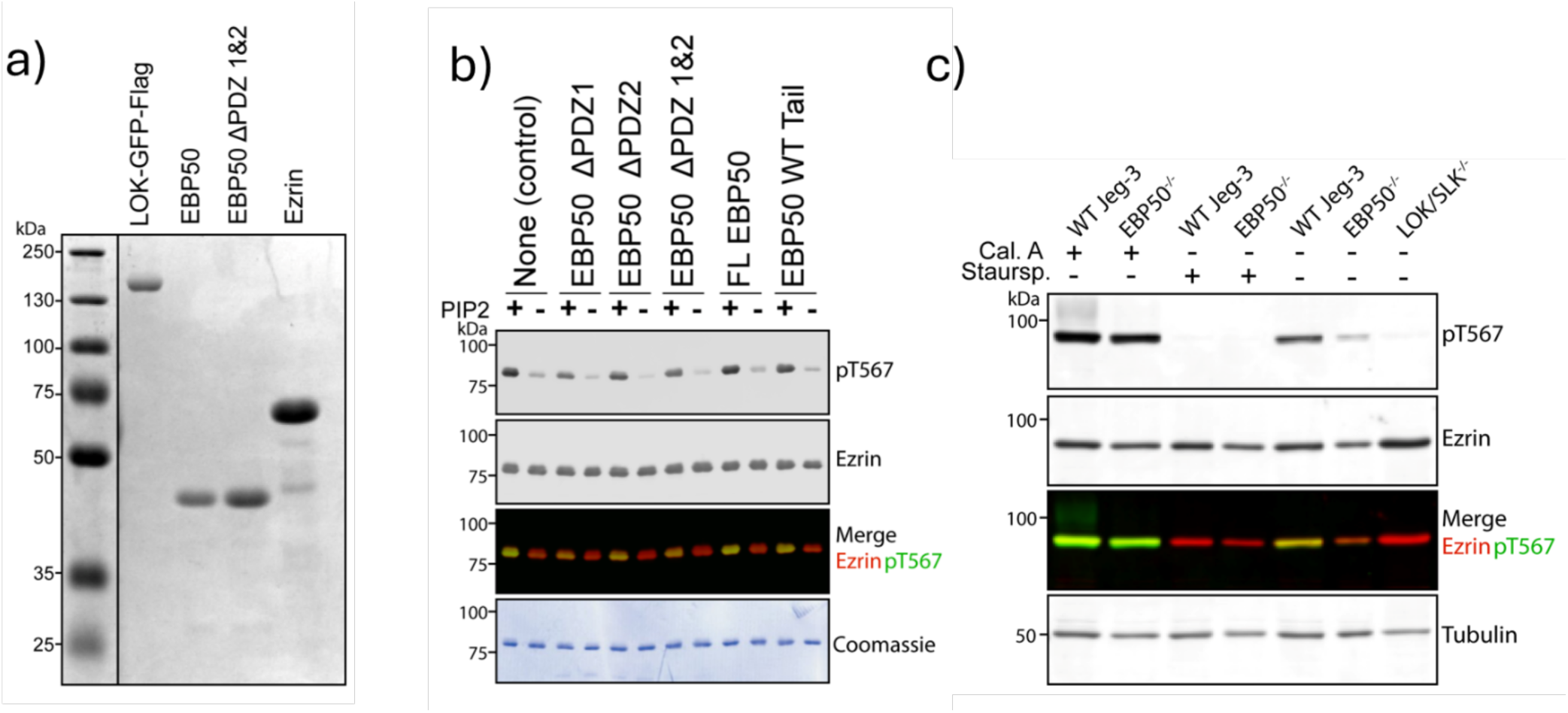
EBP50 binds to FERM after the dissociation of the CTDpT567. a) Coomassie staining of purified proteins isolated from either HEK-293T cells (LOK-GFP-Flag) or Bacterial expression (EBP50 constructs and Ezrin). b) Western blotting of biochemical in vitro kinase assays blotting for total Ezrin vs. Ezrin phosphorylated at T567 (pT567). Purified full length untagged Ezrin and LOK-GFP-Flag kinase and were added at a constant concentration 1.5mg/ml and 0.05mg/ml respectively. Prior to the addition of ATP, the proteins were incubated with 0.5mg/ml of EBP50 or EBP50 variants and PI(4,5)P_2_ was either added or withheld from duplicates of each EBP50 condition. Mutations in the two PDZ domains activate EBP50 leading to constitutively active binding between EBP50 and Ezrin’s FERM domain while EBP50’s tail domain is an unregulated Ezrin FERM binding motif. Phosphorylation of Ezrin at pT567 was PI(4,5)P_2_ dependent but independent from EBP50 binding. c) Western blotting of lysate from human WT Jeg-3 epithelial cells and Jeg-3 CRISPR knockouts of EBP50 and LOK/SLK. Cells were treated with either the phosphatase inhibitor Calyculin A (Cal. A), the kinase inhibitor Staurosporine (Staursp) or no treatment prior to lysing and blotted for total Ezrin vs. Ezrin phosphorylated at T567 (pT567). Cells lacking EBP50 have less pT567 than their WT counterparts. The effect of treatment with Cal. A or Staursp. is not altered in cells lacking EBP50 vs. WT cells indicating that EBP50 influences the turnover but not the capacity of phosphorylation/dephosphorylation of ezrin at T567. Cells lacking the endogenous ERM kinases LOK/SLK have no phosphorylation at T567.

To test if our model’s prediction that EBP50 binds subsequent to pT567 phosphorylation and competes with the ERM-CTD for binding at the FERM interface we measured pT567 levels in WT Jeg-3 human epithelial cells vs. EBP50 CRISPR KO cells produced and validated previously (41). We detected that cells lacking EBP50 had less than half the pT567 than WT Jeg-3 cells despite equal expression of total Ezrin (Figure 6c). In cells, Ezrin T567 experiences phosphate-cycling which is required for its microvillar stabilizing function (26). We sought to test if the decreased levels of pT567 observed in cells lacking EBP50 was a result of an altered capacity for Ezrin T567 to be phosphorylated or if the EBP50-FERM interactions slowed the cycling as predicted by our model. Using cell biology drug treatments of the phosphatase inhibitor Calyculin A and the broad-based kinase inhibitor staurosporine in WT vs. EBP50^-/-^ cells, we measured that WT Jeg-3 cells and cells genetically lacking EBP50 had no difference in their T567 phosphorylation or dephosphorylation capacity (Figure 6c). We concluded phosphorylation at T567 occurs before EBP50 binding to the FERM which once established, competes with the ERM-CTD for binding to the F2-F3 surface and prevents premature dephosphorylation at the T567 residue.

## DISCUSSION

The results of this study give insight into the mechanistic steps of ezrin activation upon binding to a PIP_2_-containing membrane. The microsecond-long unbiased MD simulations reveal that when FERM is anchored to the plasma membrane via the strongly negatively charged PIP_2_ phospholipid head groups, the FERM-F2 can undergo a significant conformational change, disrupting the CTD binding interface. The conformational change can be achieved at close-to-physiological PIP_2_ concentrations (4%) which are considerably lower than PIP_2_ concentrations considered in previous related studies (42). If the CTD domain is attached to the surface of the FERM F2-F3 subdomains, the energetic contribution otherwise available for FERM conformational change can facilitate the loosening of FERM-CTD interactions at the interface between F2 and the CTD. The FERM-CTD dissociation strength is then further increased by phosphorylation of T567, disrupting the extensive network of internal CTD salt bridges and reducing the strength of attachment between CTD helices H1-H2 and FERM F2 subdomain. Altogether, the unbiased atomistic MD simulations help unveil the picture of the early mechanistic steps of PIP_2_- and T567-phosphorylation-induced FERM-CTD dissociation even though their microsecond timescale is too short to sample significant opening between the FERM and CTD.

An interesting aspect of the ezrin FERM-CTD system is the perceived cooperativity of CTD association or dissociation (36). The presence of a large FERM-CTD interface area and the interconnectedness of the F2-F3 subdomains with the CTD domain via an extensive network of salt bridges means that the complete domain separation was most efficiently sampled through a coordinated, surface-wide CV via WTMetaD The difficulty in sampling the dissociation event is further exacerbated because of the inhomogeneous binding strength across the surface: CTD helix H3 is very flexible and dissociates from FERM easily, H1-H2 helices dissociate to some extent, while H4-H5 helices have remarkably strong interactions and are almost completely immobile for the entire duration of the microsecond long simulations. The H5 helix (CTD residues 576-586) is apparently immobilized due to the formation of salt bridges K211-E584, K212-E584, K577-E244 in addition to the previously suggested hydrophobic π-stacking of Phe583 with phenylalanines of the FERM F3 subdomain (not tested in this study).

The free energy profiles of FERM-CTD offer a more comprehensive understanding of what happens after FERM and CTD dissociation at PIP_2_-enriched membranes. It is found in this study that ezrin FERM-CTD system with nonphosphorylated T567 has an average ∼15 kcal/mol barrier which would correspond to 10-15 millisecond timescale according to transition state theory (43). Additionally, it is found that FERM can maintain a stable complex with bound EBP50 and has a similar FERM-EBP50 dissociation free energy profile to that of FERM-CTD-npT567. Based on a proposition from a previous study (23), EBP50 can substitute CTD in interacting with FERM at the F2-F3 surface when CTD T567 is phosphorylated and dissociates away from FERM.

Considering all the abovementioned points, we propose a pathway for FERM-CTD dissociation (Figure 7). After ezrin attachment to a PIP_2_-containing membrane, the CTD spontaneously begins to dissociate from the FERM over a millisecond-second timescale. Once CTD dissociates, the LOK N-terminal kinase domain inserts in the space between the FERM and CTD and phosphorylates CTD residue T567. The LOK kinase leaves and the CTD, liberated from FERM, gains a significant increase in configurational entropy due to the newly available degrees of freedom. Because of the high entropy-driven energetic barrier for re-association with FERM (and low barrier for escaping the partially-completed association), the CTD is unable to form a stable dissociation with FERM, and then, unrestricted by FERM, the CTD can float away to eventually reach actin filaments and tether to them. If EBP50 is available locally, the EBP50 C-terminal domain substitutes for the CTD binding to the FERM F2-F3 surface. These results are supported by the findings from *in vitro* and *in vivo* cellular biology assays which show that phosphorylation of ezrin T567 was independent of EBP50 binding. Abundant presence of phosphorylated T567 in both the wild-type and EBP50 knockout cells upon phosphatase inhibition suggests that EBP50 binds the FERM domain after CTDpT567 dissociation. The observed lower levels of pT567 in EBP50 knockout than wild-type cells in absence of phosphatase and kinase inhibitors suggest that upon EBP50 binding, CTD is more accessible for phosphate turnover as it eliminates the possibility of CTD-FERM reassociation upon dephosphorylation.

**Figure 7.**
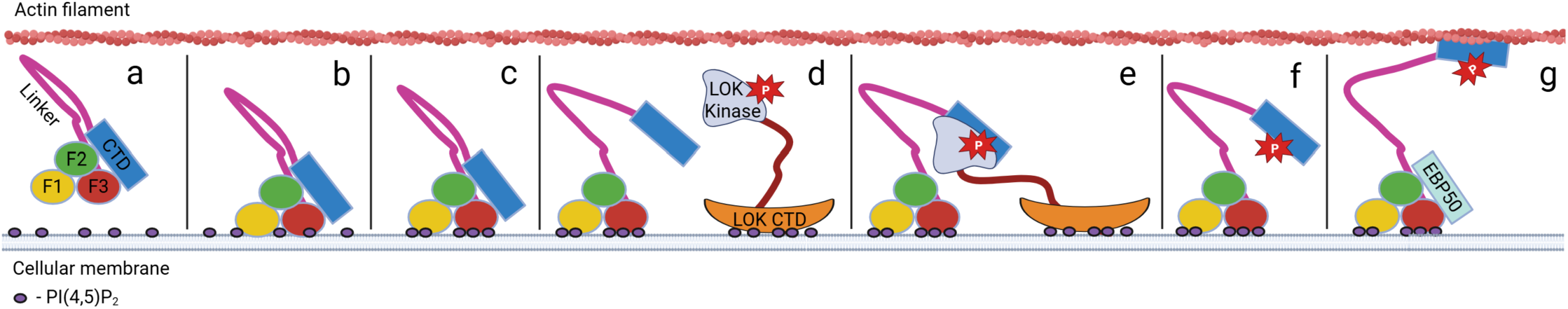
Proposed ezrin activation mechanism. a)-b) Closed state ezrin is attracted to apical cell membranes due to highly negatively charged PIP_2_ phospholipids. c) Upon attachment to membrane, FERM F1 and F3 subdomains recruit more PIP_2_. d) Ezrin wtCTD spontaneously dissociates from FERM over long timescales. e) Dissociation of CTD leaves space between FERM and CTD for LOK kinase to phosphorylate the T567 residue. f) Upon T567 phosphorylation, ezrin primarily remains in the open state due to the lower propensity to inhabit the closed state. g) CTD is enabled to float away and interact with actin filaments, thus leaving space for downstream FERM F2-F3 interactions with EBP50 or other cellular targets.

A potential avenue for future research is the explicit mechanistic analysis of the CTD T567 phosphorylation by LOK. We were able to dock ezrin CTD and LOK kinase domain (Supplementary Figure S8) and observed that the structures were stably associated throughout 500 ns solution-phase unbiased molecular dynamics runs. Despite the stable binding of the predicted structures, the exact LOK-CTD binding interfaces are yet to be determined and examined in more depth.

## CONCLUSIONS

This study helps achieve several milestones in understanding the behavior of ezrin upon binding to PIP_2_-enriched membranes. It demonstrates that upon binding to a membrane with physiological concentrations of PIP_2_ phospholipid, the ezrin FERM domain undergoes a conformational change, the energy of which can be transferred to relaxation of the attractive interactions between the CTD helices H1-H2 and FERM F2 subdomain. It is also shown that in the absence of PIP_2_, ezrin F1-F3 subdomains can recruit and anchor negatively charged DOPS residues even though aggregation of the DOPS residues does not induce conformational change as large as that induced by PIP_2_ binding. The CTD domain is found to have inhomogeneous strength of attachment to the F2-F3 subdomain surface of the FERM domain where the C-terminal end helix H5 has the strongest interactions due to multiple salt bridges and potentially hydrophobic interactions. It is also suggested that phosphorylation of T567 does not result in drastic local conformation changes but disrupts the network of CTD and F2-F3 hydrogen bonds that helps relax FERM-CTD interactions. Ultimately, our study indicates that a contact map collective variable is suitable for analyzing the thermodynamics profile of dissociation of two protein domains that are closely attached due to an extensive network of attractive interactions. A revised mechanism of ezrin activation is thus proposed which suggests that upon PIP_2_-containing membrane binding, the ezrin CTD dissociates from FERM on a millisecond-second timescale, thus allowing for T567 phosphorylation by the LOK kinase. When T567 is phosphorylated, the CTD is thermodynamically unlikely to reassociate with the FERM domain and can move away to bind to the actin filaments, which enables the FERM to interact with other cellular targets such as EBP50 at the F2-F3 subdomain surface.

## Supporting information

Supplementary Information for On the Mechanism of Ezrin Activation

## AUTHOR CONTRIBUTIONS

D.V, J.R.B, S.S.I, A.T.L, M.C.M and G.A.V conceptualized the research. D.V. and A.T.L performed simulations/experiments and analyzed results with feedback from JRB, SSI, M.C.M and G.A.V.; DV and ATL wrote the original draft. All authors contributed to the editing of the final version of the manuscript.

## ACKNOWLEDGEMENTS

This work was supported by NIGMS Grant Nos. R35GM158238 to G.A.V., R35GM156870 to A.T.L., and R01GM141372 to M.C.M. The content is solely the responsibility of the authors and does not necessarily represent the official views of the National Institutes of Health. The authors gratefully acknowledge computational resources provided by the University of Chicago Research Computing Center (https://rcc.uchicago.edu), the University of Chicago high-performance GPU-based cyberinfrastructure supported by the National Science Foundation under grant no. DMR-1828629, Frontera at the Texas Advanced Computing Center (TACC) at The University of Texas at Austin (http://www.tacc.utexas.edu) funded by the NSF (OAC-1818253), and the NIH-funded Beagle3 HPC cluster (Award Number S10OD028655). All Ezrin, LOK and EBP50 DNA constructs and CRISPR KO cell lines were a kind gift of A. Bretscher.

## DATA AND CODE AVAILABILITY

Sample simulation input files and data analysis scripts are provided in an online GitHub repository https://github.com/dovyvasi1/ezrin_activation_mechanism/. Other scripts and data are available upon request to the authors.

