## Supplementary Information for On the Mechanism of Ezrin Activation for "On the Mechanism of Ezrin Activation"

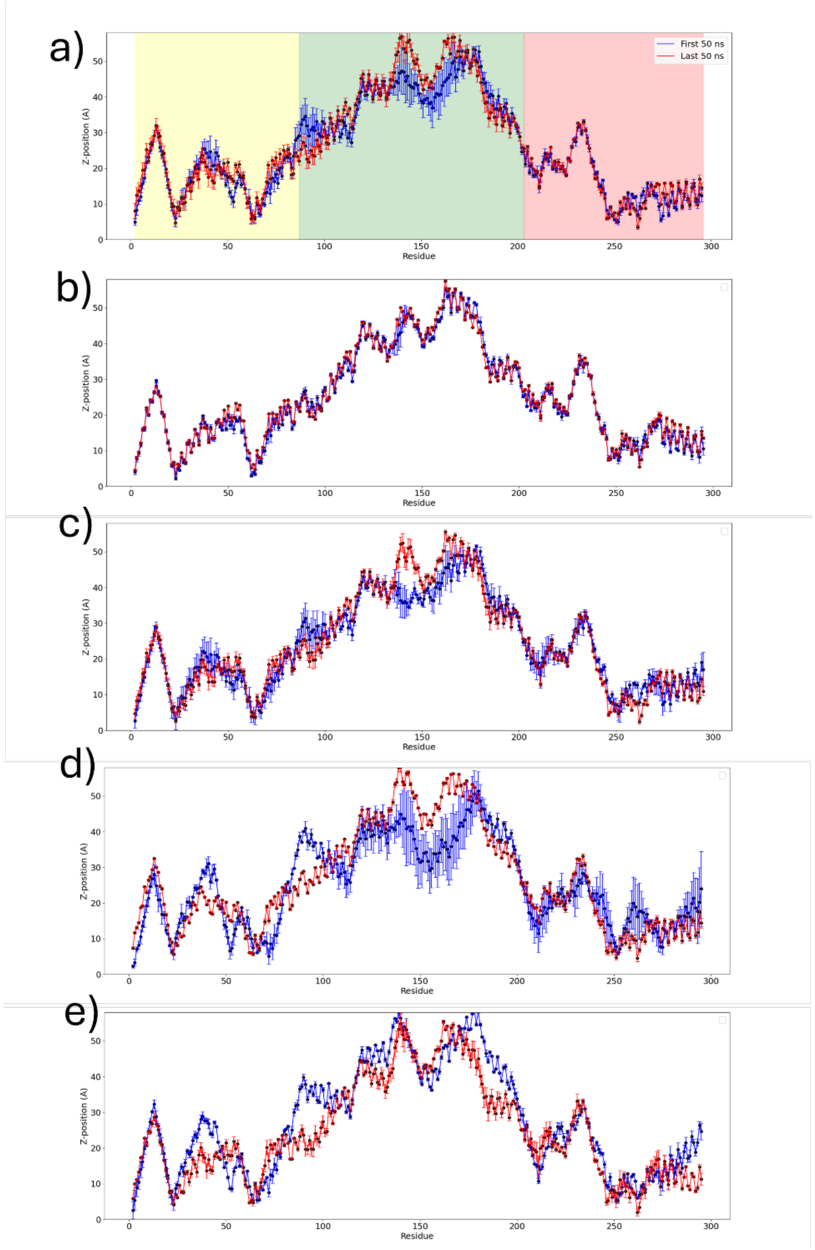


**Supplementary Figure S1.** Vertical distances of the centers of masses of FERM residues 2-296 to the mean z-position of upper membrane leaflet $PIP_2$ and DOPS head group phosphate atoms in a $PIP_2$-DOPS-DOPC membrane (images a)-c)) and mean z-distances to the upper membrane leaflet DOPS head group phosphate atoms in a DOPS-DOPC membrane (images d-e)). Red and blue plots indicate mean vertical distance of each residue in the first 50 (red) and last 50 (blue) nanoseconds of equilibrated MD simulation runs, respectively. Shaded regions correspond to FERM F1-F3 subdomains (yellow – F1, green – F2, red – F3). Images correspond to the following: a) FERM on a $PIP_2$/DOPS/DOPC membrane, b) FERM-CTD with nonphosphorylated T567 on a $PIP_2$/DOPS/DOPC membrane, c) FERM-CTD with nonphosphorylated T567 on a $PIP_2$/DOPS/DOPC membrane, d) FERM on a DOPS/DOPC membrane, e) FERM-CTD with phosphorylated T567 on a DOPS/DOPC membrane.

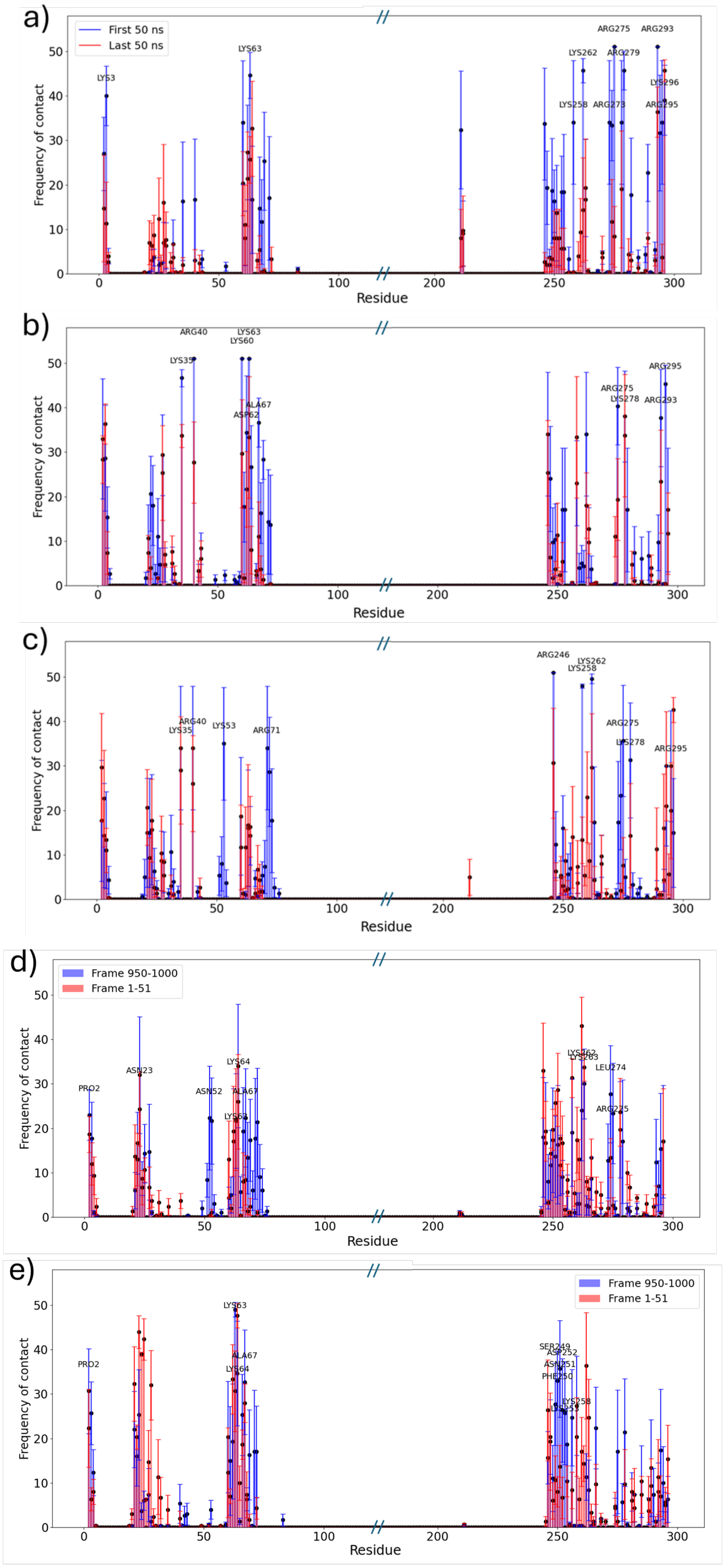
a)
First 50 ns
Last 50 ns
LYS3
LYS63
LYS262
ARG275
ARG279
ARG293
LYS296
LYS258
ARG273
ARG295
Frequency of contact
Residue
b)
ARG40
LYS35
LYS63
LYS60
ALA67
ASP62
ARG275
LYS278
ARG293
ARG295
Frequency of contact
Residue
c)
ARG40
LYS35
LYS53
ARG71
ARG246
LYS262
LYS258
ARG275
LYS278
ARG295
Frequency of contact
Residue
d)
Frame 950-1000
Frame 1-51
PRO2
ASN23
ASN52
LYS64
ALA67
LYS62
LYS262
LYS263
LEU274
ARG275
Frequency of contact
Residue
e)
Frame 950-1000
Frame 1-51
PRO2
LYS63
ALA67
LYS64
SER249
ASP252
ASN251
PHE250
LYS258
Frequency of contact
Residue

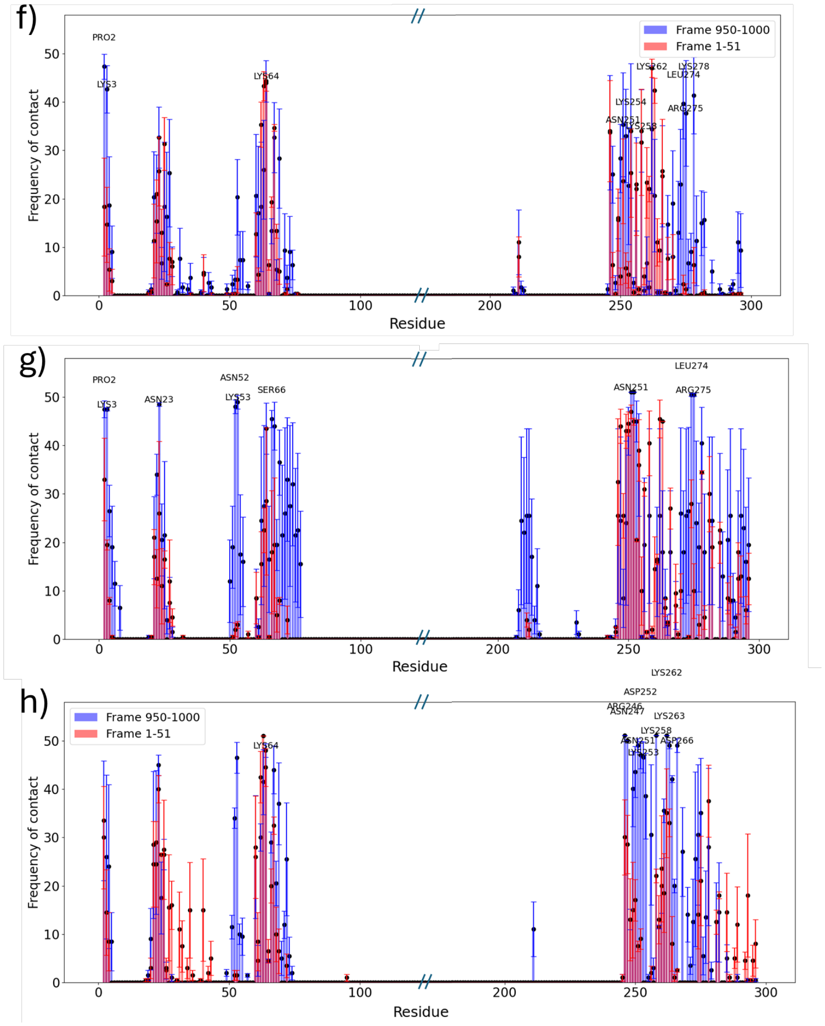


**Supplementary Figure S2. FERM F1 and F3 lobes attract negatively-charged phospholipid head groups, $PIP_2$ displaces less-charged DOPS head groups at the FERM F1-F3 surface.** Frequency of less-than-10 Å contacts between FERM residues and $PIP_2$ head group (a-c) phosphate atoms as well as FERM residues and DOPS head group (d-h) phosphate atoms in the first 50 (red) and last 50 (blue) nanoseconds of equilibrated MD simulation runs. a) FERM at a $PIP_2$/DOPS/DOPC membrane, b) FERM-CTD with nonphosphorylated T567 at a $PIP_2$/DOPS/DOPC membrane, c) FERM-CTD with phosphorylated T567 at a $PIP_2$/DOPS/DOPC membrane, d) FERM and a $PIP_2$/DOPS/DOPC membrane system, e) FERM-CTD npT567 and a $PIP_2$/DOPS/DOPC membrane system, f) FERM-CTD pT567 and a $PIP_2$/DOPS/DOPC membrane system, g) system of FERM and a aDOPS/DOPC membrane, and h) system of FERM-CTD pT567 and a DOPS/DOPC membrane. The labeled residues are the residues that had the highest frequency of contact with $PIP_2$ head group phosphate after 1000 ns unbiased MD simulation runs. As can be seen, positively charged Arg, Lys residues established most stable contacts with $PIP_2$ head groups.

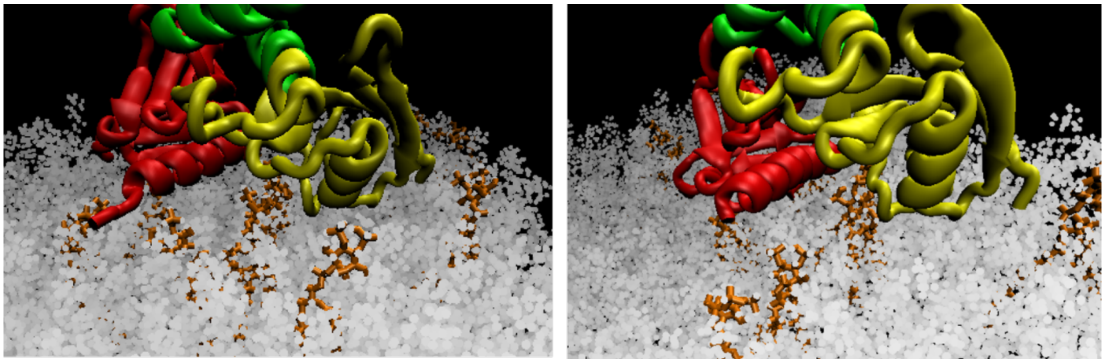

**Supplementary Figure S3.** $PIP_2$ phospholipds (orange) become localized in the cleft between F2 (yellow) and F3 (red) subdomains of FERM domain and thus affect the local conformation in the F3 subdomain, particularly in helix that contains residues 273-296. Snapshots of two replicas of unbiased MD simulation of FERM domain at a $PIP_2$/DOPS/DOPC membrane at t=1000 ns.

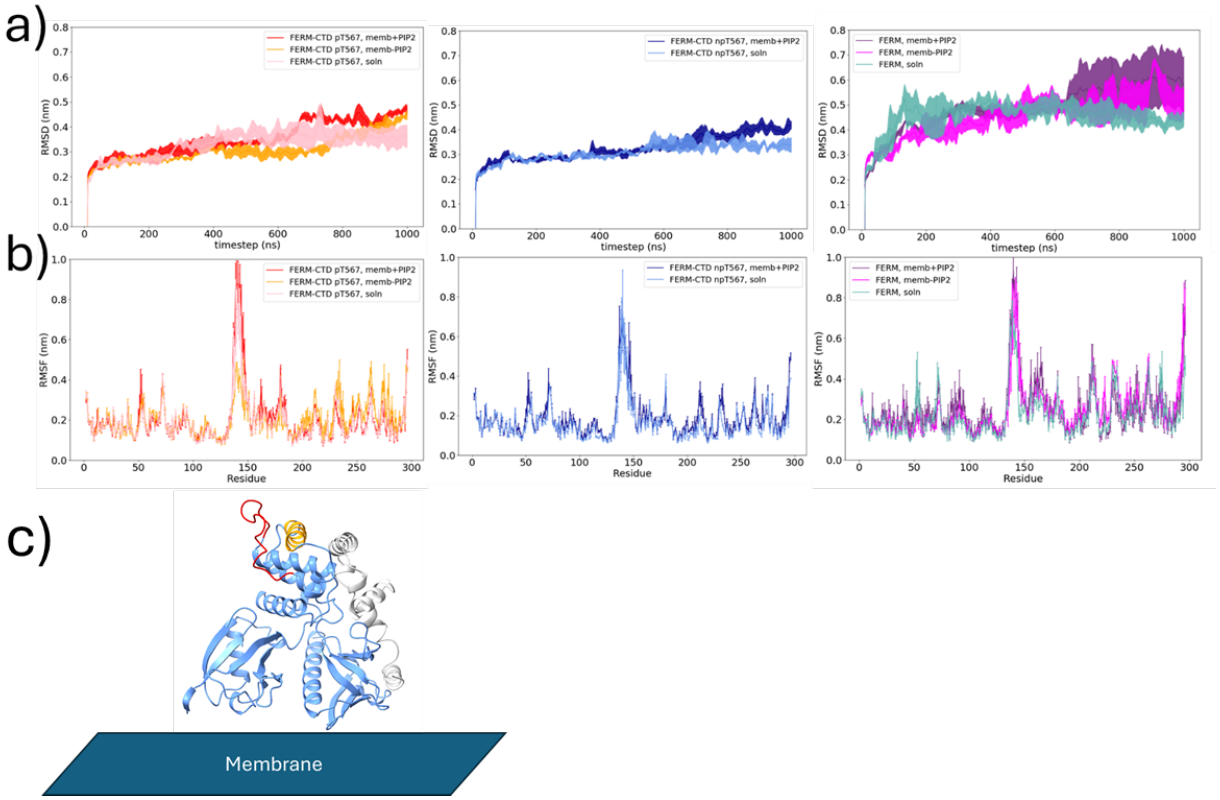


**Supplementary Figure S4.** Structural stability analysis of the FERM domain (ezrin residues AA2-AA296) in different ezrin systems a) Root mean square displacement values (RMSD) values throughout the course of the 1 μs unbiased MD simulations, b) root mean square fluctuation values (RMSF) during the last 100 ns of the 1 μs unbiased MD simulations. FERM domains without an attached ezrin CTD domain feature much larger RMSD values throughout the course of the simulation. The rate of conformational change increase is lower in FERM-CTD systems potentially because CTD constrains the motion of the FERM F2 subdomain. Notation: pT567 – system with phosphorylated T567, npT567 – system with nonphosphorylated T567, memb+$PIP_2$ – system includes a $PIP_2$/DOPS/DOPC membrane, memb–$PIP_2$ – system includes a DOPS/DOPC membrane, soln – simulation conducted in a 0.15 M KCl solution. c) The initial structure of FERM with highlighted residue groups AA136-AA152 (red) and AA167-AA179 (orange) which feature substantial fluctuations.

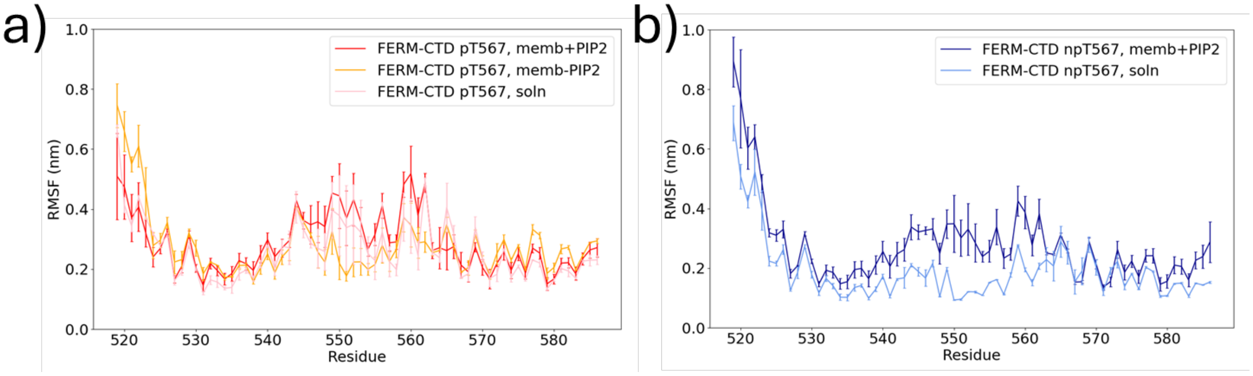


**Supplementary Figure S5.** Root mean square fluctuation (RMSF) analysis for CTD residues of systems with (a) phosphorylated and (b) nonphosphorylated T567.

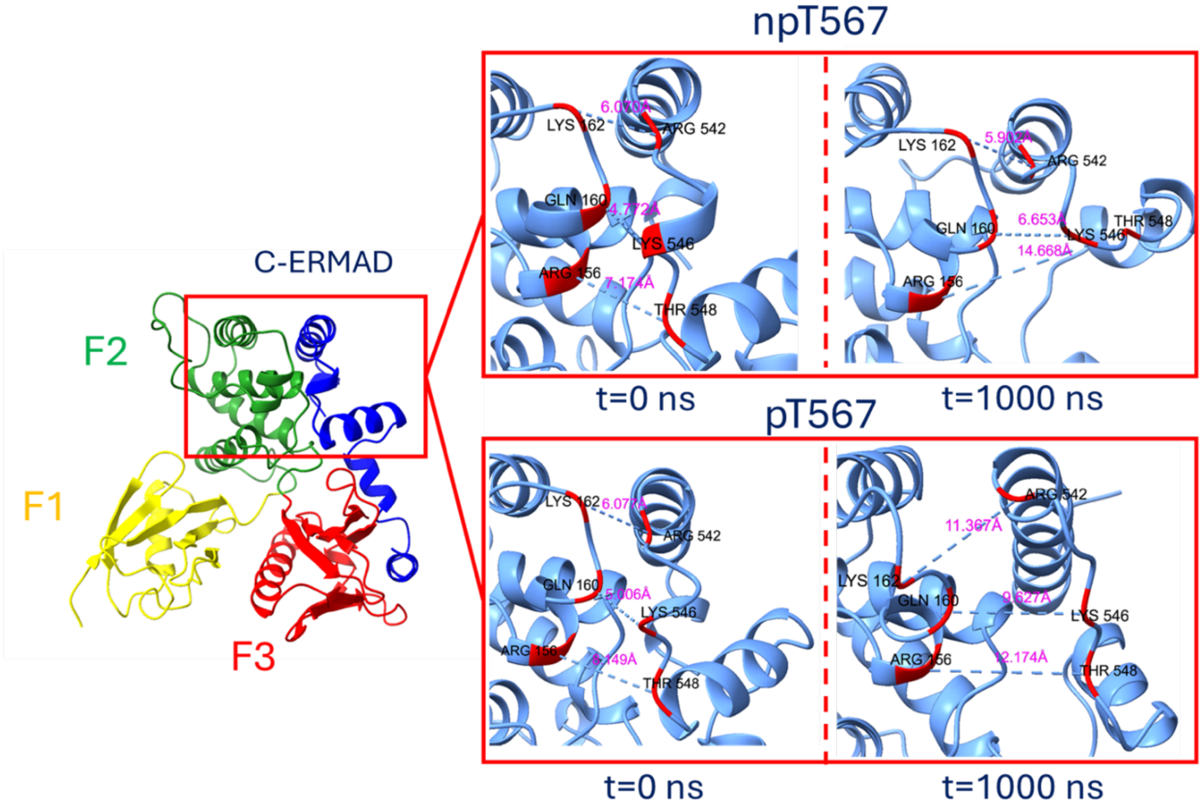


**Supplementary Figure S6.** Significant opening between CTD residues AA538-AA546 and FERM F2 lobe is only observed in the FERM-CTD system with phosphorylated T567 at a $PIP_2$/DOPS/DOPC membrane. Left – crystal structure of ezrin FERM and C-ERMAD (PDB ID:4RM9). Right – distance between alpha carbon atoms of residue pairs K162-R542, Q160-K546, R156-T548 in the beginning of the equilibrated unbiased MD simulations (t=0 ns) and the end of the 1 μs simulation (t=1000 ns) for versions of the system that contain either a nonphosphorylated T567 (npT567, top) or phosphorylated T567 (pT567, bottom).

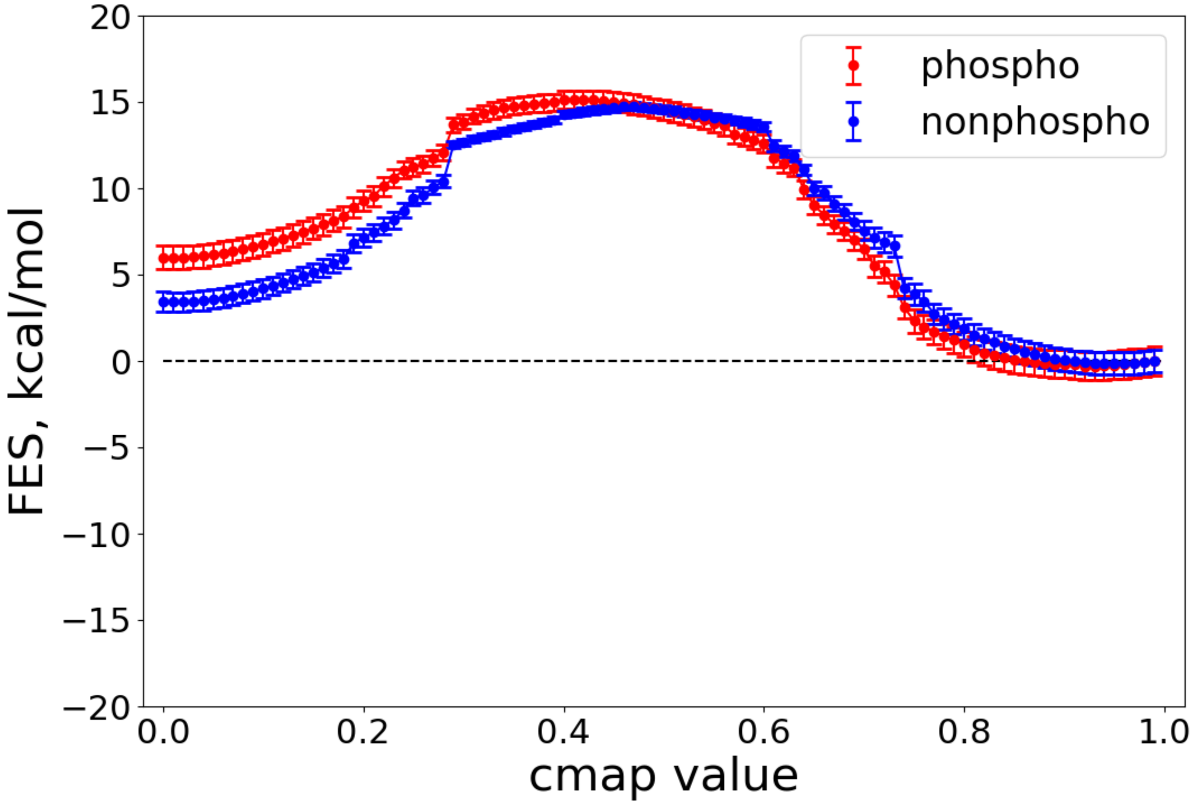


**Supplementary Figure 7.** Free energy profiles for solution-phase FERM-CTD systems with wtCTD (blue) and pT567CTD (red) obtained from contact map well-tempered metadynamics. The dissociation barriers were 14.9 ± 0.2 kcal/mol for wtCTD and 15.4 ± 0.5 kcal/mol for pT567CTD, and reassociation barriers were 11.4 kcal/mol for wtCTD and 9.4 kcal/mol for pT567CTD. The sharper drops/increases at contact map values of about 0.3 and 0.7 correspond to sharper drops at those collective variable values observed in separate reweighted free energy profiles.

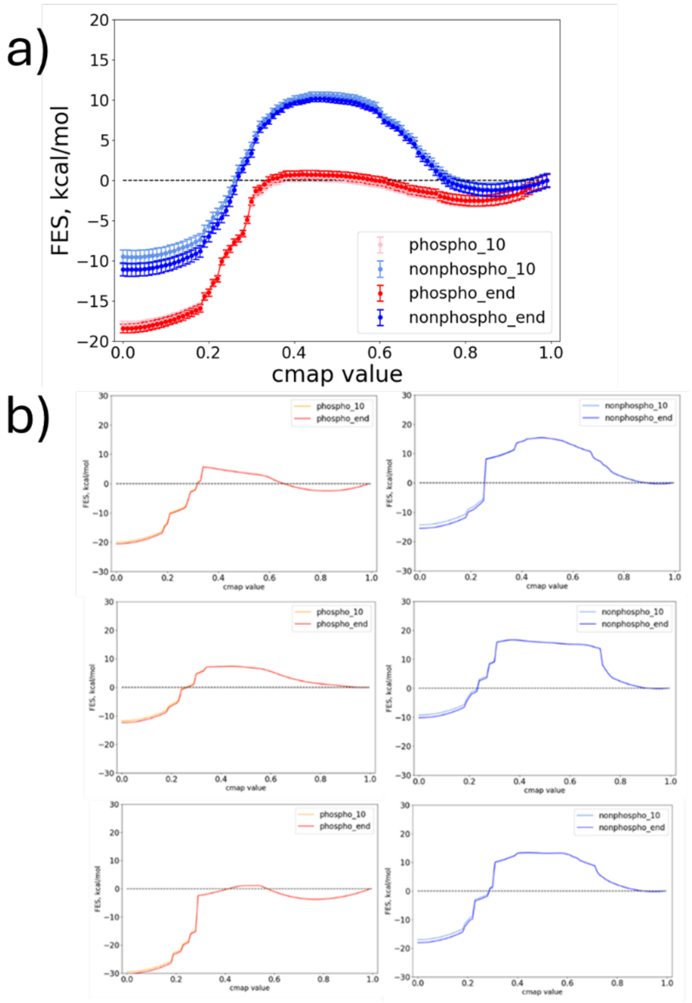


**Supplementary Figure S8.** a) Convergence of the WTMetaD runs. Convergence is observed from the similarity of the free energy surfaces at the end of metadynamics runs (phospho_end and nonphospho_end) and the surfaces obtained without the last 10 ns of metadynamics data (nonphospho_10 and phospho_10). b) Representative individual reweighted contact map metadynamics free energy surfaces for phosphorylated (left) and nonphosphorylated (right) systems. The convergence of the calculation is observed from the negligible difference between the final FES curves (phospho_end, nonphospho_end) and the FES curves obtained without the last 10 ns of metadynamics data (phospho_10 and nonphospho_end).

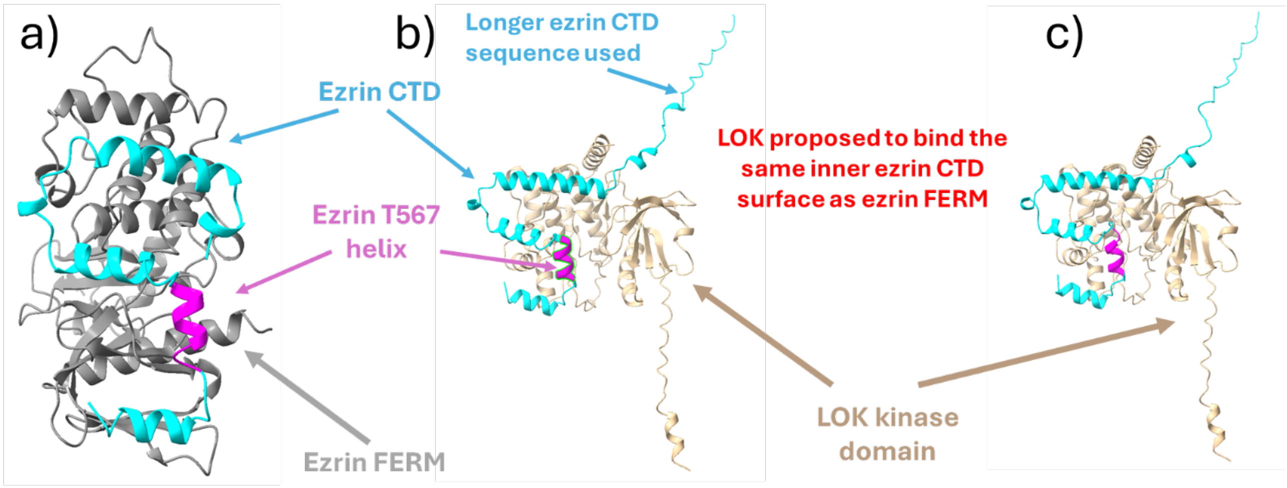


**Supplementary Figure S9.** Ezrin CTD domain and LOK kinase domain docking results. The docking procedure contained 3 major steps: generation of highly likely ezrin CTD-LOK kinase with AlphaFold 3.0[14], selection of the structures that had substantial contacts between T567 and kinase domain residues, and verification of binding with ClusPro software. a) FERM-CTD crystal structure with highlighted T567-containing H4 helix, b) AlphaFold3 predicted association of a longer-sequence CTD with the LOK kinase domain, c) highest-fidelity ClusPro docking result for the AlphaFold3-predicted CTD and kinase domain structures.

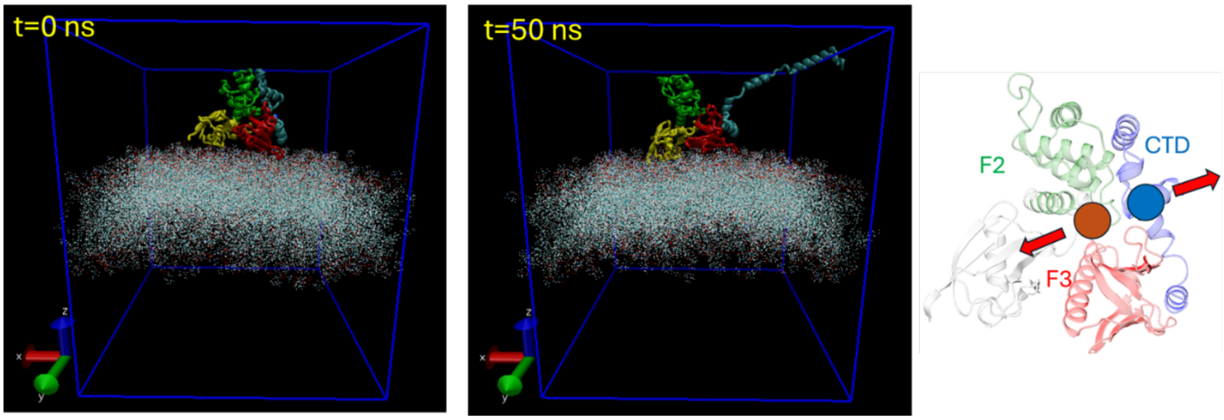


**Supplementary Figure S10.** Snapshots from well-tempered metadynamics simulation runs where the collective variable is defined as the distance between the centers of mass of F2-F3 subdomains and the center of mass of the C-ERMAD domain. Well-tempered metadynamics simulations with distance-based CVs were not able to provide a full dissociation of CTD domain from the FERM domain. Simple CVs (distance between the centers of masses of CTD and F2-F3) and more complex CVs alike (pathCV where path variable is defined from the distance of center of mass (COM) of F2 to the COM of its near portion of CTD and the distance of COM of F3 to the COM of its near portion of CTD) were unable to sample the dissociation of the entire CTD domain from the FERM domain. In agreement to unbiased MD simulation results which showed that F3-CTD association was much stronger than F2-CTD association, these metadynamics runs only showed dissociation of CTD from F2 while F3-CTD remained tightly attached throughout the course of tens- and hundreds of nanoseconds of simulations. Similar outcomes were observed in steered MD enhanced sampling simulations where only F2 and CTD were able to separate. While we were not able to fully probe the dissociation with these methods, we do note the common mechanism showing CTD unbinding from the F2 domain first, consistent with our previous suggestion that the CTD is mostly weakly attached there.

**Strengths, limitations, and future directions**

In this study, we investigated the microsecond-scale ezrin FERM-CTD interactions upon FERM binding to $PIP_2$ and the free energy profiles of full FERM-CTD dissociation for systems with phosphorylated and nonphosphorylated T567 using one-dimensional contact map WTMetaD simulations. Our results give additional support that $PIP_2$ anchoring is crucial for the phosphorylation of T567 and indicate that phosphorylation of T567 is independent of EBP50 binding to FERM domain. Combined, computational and experimental results provide new understanding of ezrin FERM-CTD dissociation mechanism in which LOK kinase phosphorylates T567 after FERM-CTD dissociation, substantially reduces the affinity of CTD to FERM and consequently leaves space for downstream events such as the EBP50-FERM association.

It is important to note that there are certain drawbacks too. Although early steps of ezrin activation are observed in unbiased atomistic simulations, the system size and computational constraints prevent a complete atomistic investigation of what happens afterwards, including an analysis of explicit LOK kinase phosphorylation of the T567 residue. A possible bias to the observed WTMetaD results also comes from the exclusion of the ~200 residue ezrin linker domain, even though the linker region connection to CTD is relatively loose (results not shown) and the main contributor to strong FERM-CTD attachment strength is the network of the salt bridges between FERM subdomains F2-F3 and CTD. The WTMetaD results might have been biased to some degree by the high relative speed of contact map CV coordinate crossing which caused some sample systems get occasionally stuck in either the closed or the open states in the later stages of the WTMetaD simulation (results not shown). Some bias could have been introduced from the selection of the initial WTMetaD frame selection system which were averaged to get the final free energy profiles; however, it is unlikely that the profiles would be substantially different for different sets of starting frames because FERM-CTD and FERM-membrane association does not change substantially throughout the course of the 1 μs simulations.

It should also be emphasized that the FERM-CTD system used in this study is a working model that disregards the effects of the ~200 residue linker region. The results of FERM-CTD dissociation thermodynamics obtained from using this model are deemed transferable to the free energy profiles of the full-length ezrin because the linker region connection to CTD is fairly loose (results not shown) and the main contributor to strong FERM-CTD attachment strength is the network of the salt bridges between FERM subdomains F2-F3 and CTD.
